# Examining population structure across multiple collections of Cannabis

**DOI:** 10.1101/2022.07.09.499013

**Authors:** Anna Halpin-McCormick, Karolina Heyduk, Michael B. Kantar, Nicholas L. Batora, Rishi R. Masalia, Kerin Law, Eleanor J. Kuntz

## Abstract

Population structure of *Cannabis sativa* L. was explored across nine independent collections that each contained a unique sampling of varieties. Hierarchical Clustering of Principal Components (HCPC) identified a range of three to seven genetic clusters across datasets with inconsistent structure based on use type indicating the importance of sampling particularly when there is limited passport data. There was broader genetic diversity in modern cultivars relative to landraces. Further, in a subset of geo-referenced landrace accessions, population structure was observed based on geography. The inconsistent structure across different collections shows the complexity within *Cannabis*, and the importance of understanding any particular collection which could then be leveraged in breeding programs for future crop improvement.

## Introduction

*Cannabis sativa L.* is an annual flowering herb which has been domesticated multiple times for food, fiber and medicine over the last twelve thousand years (Hillig, 2005; Clarke & Merlin, 2013; Clarke & Merlin, 2016; Ren et al., 2021). *Cannabis* is popularly known for its psychoactive effects; however, it is its medicinal capacity is driving increased production (Punja & Holmes 2020). The compounds tetrahydrocannabinol (THC) and cannabidiol (CBD) are the most studied due to their potential in pain management (Walker & Huang, 2002; Alexander, 2020; Bicket et al., 2023), as a multiple sclerosis treatment (Svendsen et al., 2004), for epilepsy management (Charlotte’s Web (CW2A) US Plant Patent No. PP30,639 P2; Perucca, 2017), for reduction in nausea (Parker et al., 2011) and as an appetite stimulant (Badowski & Perez, 2016). Today, *Cannabis* is broadly divided into non-drug and drug-type cultivars (**Table 1**).

**Table 1.** Definitions related to the different types of germplasm that were used in this study.

| Type | Class | Definition |
| --- | --- | --- |
| Feral | Used as both Non-drug and Drug type | Plants that have escaped cultivation and are now growing in the wild without human intervention. These accessions have no influence of human selection. |
| Landrace | Used as both Non-drug and Drug type | Cultivars are introduced to a region by humans and then become locally adapted to a specific geography over time mostly through indirect selection by farmers and natural selection. |
| Modern | Used as both Non-drug and Drug type | Cultivars that have been intentionally bred and selected by humans using advanced breeding techniques (genetics and statistics) with the goal of enhancing specific traits. These cultivars have been developed in recent years or decades and may not have the same regional or historical ties as landrace strains. |
| Hemp | This material is used for fiber - Non-drug type | Samples that had names of hemp used for grain and fiber or wild collected feral plants (no chemical analysis to confirm hemp or marijuana) |
| Type III | Non-drug type | (CBD-dominant): cannabis flower defined as hemp in the U.S with <0.3% THC with a wide range of CBD (average 12% 30:1 CBD:THC) |
| Type II | Drug-type | (CBD:THC): balanced ratio of THC:CBD (1:1) |
| Type I | Drug-type | (THC-dominant): modern cannabis strains found in the legal U.S. medical and adult use market (generally >10% THC, average 21% THC, usually 30:1 THC:CBD ratio) |

Due to the classification of *Cannabis* as a Schedule I narcotic in the United States, research during the 20^th^ century was largely restricted (Hurgobin et al., 2021). However, the 2018 United States Farm Bill reduced these restrictions, with many states now having reduced regulations (Mead, 2019). In 2020 the Drug Enforcement Administration expanded research licenses (Ryan et al., 2021) leading to increased *Cannabis* research. However, due to past restrictions C*annabis* has not fully benefited from scientific tool developments (e.g. molecular marker tools, heterotic pattern development) of the last century. Further, drug control laws and prohibition have constrained formal documentation often resulting in unverifiable and anecdotal cultivar origins (Duvall, 2016). However, there has been recent work to develop tools and initiate breeding (Toth et al., 2020; Petit et al., 2021; Woods et al., 2021; Toth et al., 2022; Woods et al., 2023).

While there is still some debate, the taxonomy of *Cannabis* has moved towards a monotypic description of the genus (McPartland 2018; McPartland & Small 2020). *Cannabis* populations have been partitioned using different methods (e.g., genetic, chemical and phenotype) with different populations showing different patterns (de Meijer et al., 2003; Lynch et al., 2016). There are also examples of studies using regional, ecotype, and use-type to understand population partitions (Soorni et al., 2017; Zhang et al., 2020; Carlson et al., 2021; Ren et al., 2021). These studies have found contrasting results due to contested definitions and different samples. In addition to understanding species and population delineation, previous genetic work has explored the cannabinoid metabolic pathways (Guerriero et al. 2017; Guerriero et al., 2019; Allen et al., 2019; McKernan et al., 2020; van Velzen & Schranz 2021).

Understanding population structure provides insight into evolutionary relationships and facilitates the identification of cultivars that have value for breeding practices. Further, understanding genetic relationships can help reconstruct pedigrees and genetic relationships which have been lost due to a century of prohibition. Clarification of cultivar relationships could provide more concrete reproducible results in addition to the ethnohistorical information and spoken accounts that underpin current research. The molecular genetic profiles of *Cannabis* cultivars will enhance our understanding of them, providing a valuable tool to confirm marketing claims independently of relying solely on visual characteristics. This, in turn, can contribute to a more reliable and sustainable industry.

Previous work has used a range of sequencing methods, reference genomes, and sampling schemes (Small & Cronquist, 1976; Clarke, 1987; van Bakel et al., 2011; Duvall 2016; Soorni et al. 2017; Soler et al., 2017; Maoz 2020; Hurgobin et al. 2021; Grassa et al. 2021). In an attempt to understand previous studies (eight publicly available datasets) as well as a newly generated dataset we used a common single nucleotide polymorphism (SNP) calling pipeline and the same reference genome (Grassa et al., 2021) to explore population structure present across different germplasm collections and to identify potential samples that can be explored as the basis for breeding.

## Materials and Methods

### Sequence Data Acquisition

Raw sequence data from Soorni et al. 2017 (PRJNA419020), Lynch et al., 2016 (PRJNA317659), Phylos Biosciences (PRJNA347566 & PRJNA510566), Courtagen Life Sciences (PRJNA297710), and Sunrise Genetics (PRJNA350539) were downloaded from National Center for Biotechnology Information (NCBI: https://www.ncbi.nlm.nih.gov/) using the SRA toolkit (https://hpc.nih.gov/apps/sratoolkit.html). Data from Medicinal Genomics, where 61 paired samples were provided for bulk download (Medicinal Genomics 61 - https://www.medicinalgenomics.com/kannapedia-fastq/) and an additional 289 samples (Medicinal Genomics StrainSEEK v1) were individually downloaded from each cultivar page. The last data source used here was developed by LeafWorks Inc., consisting of 498 individuals. Full dataset descriptions are available in **Table 2**.

**Table 2.**
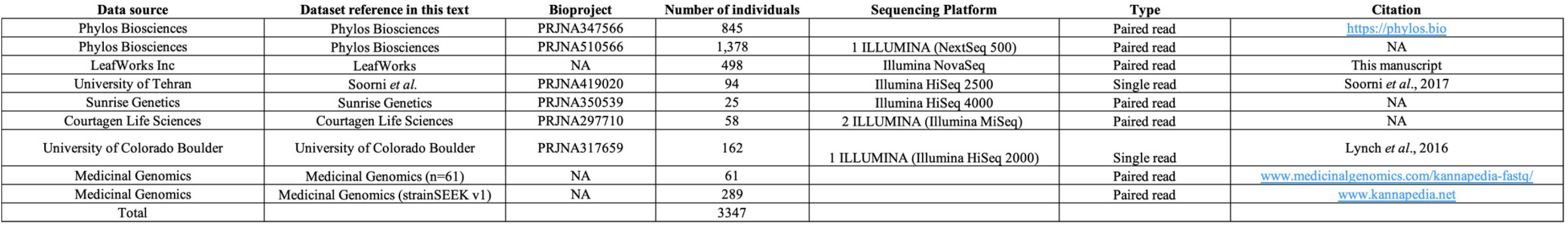
Data sources used for this project. Light grey indicates other public datasets which were not utilized in this study.

### Sample Name Acquisition

Sample names were assigned to individual samples as supplied by authors in supplemental materials of publication or through the metadata supplied through NCBI. All individual line assignments can be seen in **Tables S1-S9**. For Phylos Biosciences datasets, each SRR number was searched in NCBI in the SRA database. For the n=845 and the n=1,378 datasets this facilitated the association of SRR numbers with cultivar names from the “Sample” section and aided in matching the sample to the genotype information sheet on the Phylos Biosciences website (https://phylos.bio/). The links to the matching genotype report page for each SRA sample have been included in the metadata of the supplemental tables (N=845 **Table S2** and n=1,378 **Table S3**).

### Use-type Category Assignment

Different meta-data for each dataset was used to identify the use-type (**Table 1**). For Phylos Biosciences (https://phylos.bio/) and Medicinal Genomics (https://www.kannapedia.net). For the Medicinal Genomics dataset, where no information was reported in the “Plant Type” section on individual strain pages, the cannabinoid section on strain pages which reports percentages of THC and CBD as well as other cannabinoids was used to assign type to individual samples, with well-known hemp variety names facilitated by the EU Plant variety database https://ec.europa.eu/ (eg. Santhica, Carmagnola, Fedora, Felina). For the LeafWorks Inc. dataset, type associations were provided for 101 landrace samples and 44 hemp samples with remaining use-type associations assigned through searching sample names on https://www.leafly.com or https://www.wikileaf.com. For Soorni et al. 2017 dataset a recent publication used chemistry of these same accessions to determine use-type (Mostafaei Dehnavi et al. 2022). For the remaining datasets (Sunrise Genetics, Lynch et al., 2016, and Courtagen Life Sciences) sample names were searched on https://www.leafly.com or https://www.wikileaf.com for assignment to a category of use-type (**Tables S1-S9**). Use-types were not evenly represented and this uneven representation of different use-types may influence conclusions related to the genus overall (**Tables S1-S9**).

### Sequence Data Processing

Where demultiplexing was required, barcodes were acquired from the supplemental materials and removed using the software SABRE (version 1.0 - https://github.com/najoshi/sabre). All dataset fastq files were checked for adapter sequence content using the FASTQC (version 0.11.8- Andrews, 2010). Datasets were examined post FASTQC using MULTIQC (Ewels et al., 2016). Where adapters were present, TRIMMOMATIC (version 0.39 - Bolger et al., 2014) was used to remove these sequence elements. The software SKEWER (Jiang et al. 2014) was used to trim adaptors from Phylos Biosciences n=1,378 dataset. Some data from Lynch et al. 2016 (PRJNA310948) was also not included as PRJNA310948 appears to contain duplicates of samples from PRJNA317659 both released in 2016. Therefore, only PRJNA317659 was used. The Medicinal Genomics’ Kannapedia site contains samples that have been sequenced across a variety of platforms, for consistency here we used the samples from StrainSEEK v1 (n=289).

Reads were then aligned to the CBDRx genome (Grassa et al. 2021) using BWA-MEM (version 0.7.17 - Li, 2013). SAMTOOLS (version 1.9 - Li et al., 2009) was used to convert SAM files to BAM files and mapped reads were sorted for a mapping quality of 30 or above. BCFTOOLS (version 1.9 - Danecek & McCarthy, 2017) using the mpileup function was used to generate SNPs and create VCF files. Samples were filtered using VCFtools (version 0.1.16 - Danecek et al., 2011) for a minor allele frequency of 0.05, Hardy-Weinberg Equilibrium (0.05), and a maximum missingness of 10%. After filtering, data were analyzed using the SNPRelate (Zheng et al. 2012), FactoMineR (Lê et al., 2008) and factoextra (Kassambara & Mundt, 2017) packages in RStudio (version 1.4.1106 - R Core Team, 2013).

### Nucleotide Diversity Calculation

VCF files for known modern cultivars and landraces were separately merged into a single VCF file. Nucleotide diversity (*π*) was calculated using VCFtools with a 10,000 bp sliding window across the strictly filtered files for each dataset. Changes in *π* across chromosomes were plotted in RStudio using the ggplot package (Wickham, 2011).

### Population Structure and Phylogenetic Analysis

VCFtools was used to generate MAP and PED files. These were then used to generate BED, BIM, and FAM files in the software PLINK (version 1.9 - Purcell et al., 2007). For each dataset population structure across a range of population partitions was assessed in fastSTRUCTURE (version 1.0 - Raj et al., 2014). The optimal number of K was also examined for each dataset using the elbow and silhouette methods in the FactoMineR and factoextra packages. In addition, each dataset was examined using Principal Component Analysis (PCA) in SNPRelate (Zheng et al., 2012). Only bi-allelic SNPs further filtered for linkage disequilibrium (0.2) were used for the PCA and Hierarchical Clustering on Principal Components (HCPC) (**Fig. 1-2 and Fig. S2-6**). A Maximum Likelihood (ML) phylogenetic tree was constructed for the LeafWorks Inc. dataset, a VCF file was converted to NEXUS and FASTA format using the software package VCF2PHYLIP (version 2.6 - https://github.com/edgardomortiz/vcf2phylip). Ambiguities were changed to “N” where observed. Multiple sequence alignment was performed using MAFFT (version 7.475- Katoh & Standley 2013) and this was submitted to the software ModelTest-NG (version 0.1.6 - Darriba et al. 2020) to best evaluate the substitution model to be used.

**Fig. 1.**
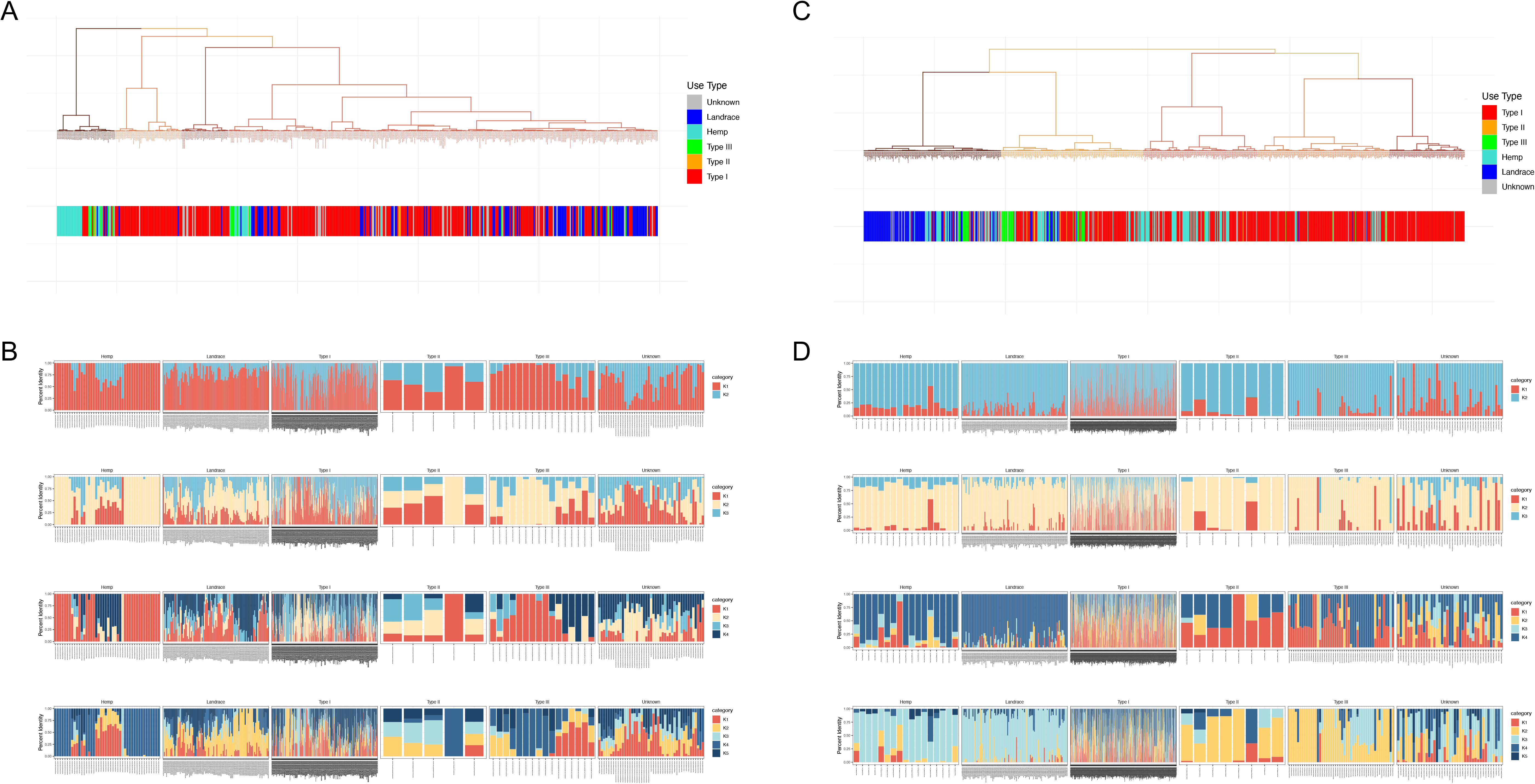
Examining hierarchical clustering on principal components (HCPC) and population structure in the LeafWorks Inc. (n=498) and Phylos Biosciences (n=845) datasets. In each case population genetic clustering was conducted based only on nuclear genetic SNPs while reported use-type within the dataset is below in solid bars to facilitate interpretation based upon community standards **(A)** Hierarchical cluster dendrogram from 520 nuclear SNPs for the LeafWorks Inc. dataset with use-type indicated below. Use-type are pictured below (Type I=288, Type II =5, Type III=16, Hemp=44, Landrace=101 and Unknown=44) **(B)** Visualization of population structure and admixture from 1,405 nuclear SNPs for the LeafWorks Inc. dataset using the fastSTRUCTURE software (k=2-5) with the optimal number of K being 4 using the silhouette method **(Fig. S9-10) (C)** Hierarchical cluster dendrogram from 292 nuclear SNPs for the Phylos Biosciences dataset with use-type indicated below. Use-type accessions include Type I=479, Type II=8, Type III=46, Landrace=127, Hemp=143 and Unknown=42 **(D)** Visualization of population structure and admixture from 385 nuclear SNPs for the Phylos Biosciences dataset using the fastSTRUCTURE software (k=2-5) with the optimal number of K being 3 using the Silhouette method **(Fig. S9-10)**.

**Fig. 2.**
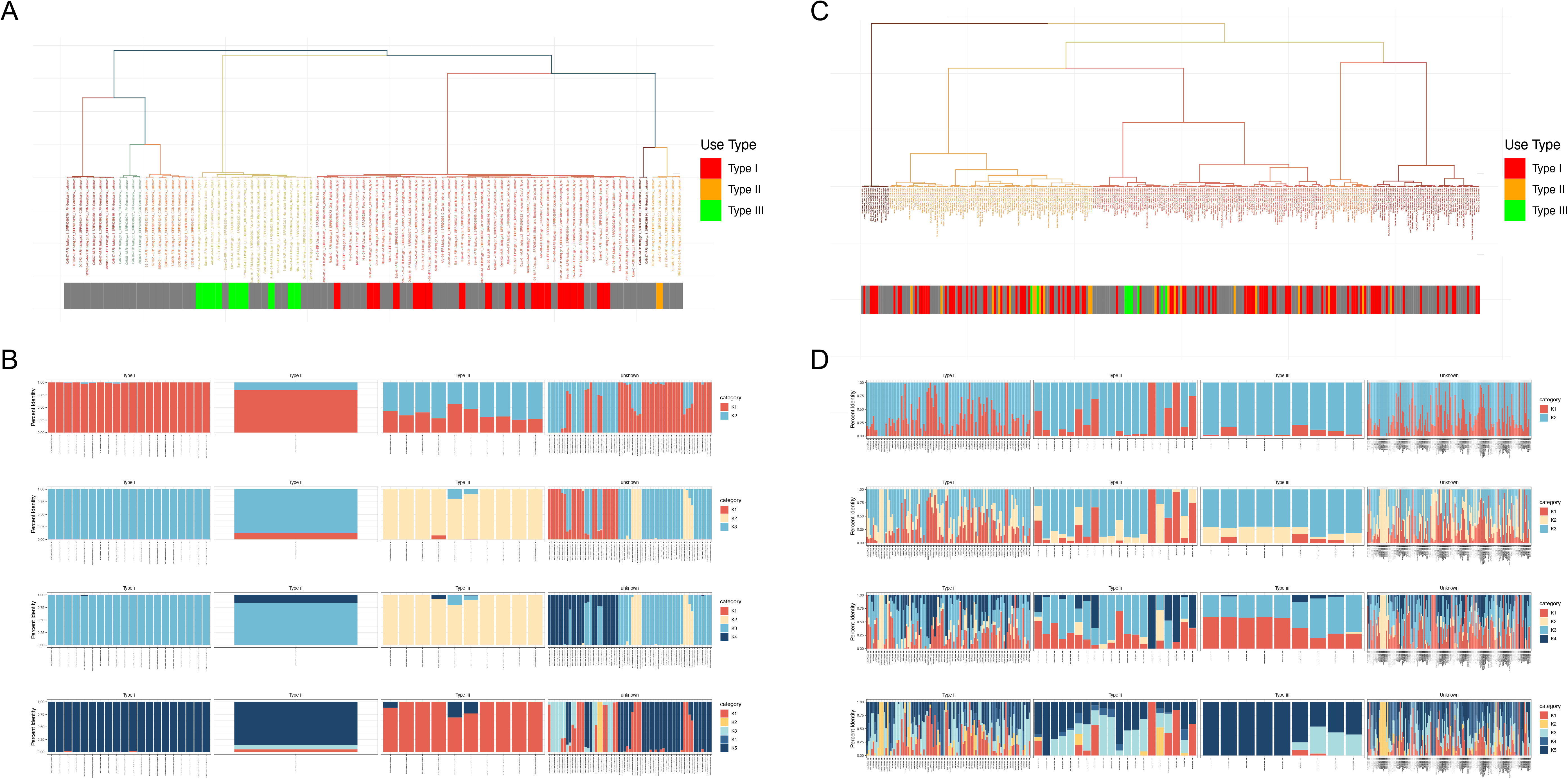
Examining hierarchical clustering and population structure in the Soorni et al. 2017 (n=94) and the Medicinal Genomics StrainSEEK V1 (n=289) datasets. In each case clustering was conducted based on nuclear genetic SNPs while reported use-type within the dataset is below in solid bars to facilitate interpretation based upon community standards **(A)** Hierarchical cluster dendrogram from 6,865 nuclear SNPs for the Soorni et al. 2017 dataset with use-type of each accession indicated below. Use-type are pictured below (Type I=20, Type III=10, Type II=1, Landrace=78 and Unknown=63) **(B)** Visualization of population structure and admixture from 33,629 nuclear SNPs for the Soorni et al. 2017 dataset using the fastSTRUCTURE software (k=2-5) with the optimal number of K being 3 using the silhouette method **(Fig. S9-10)** (**C)** Hierarchical cluster dendrogram from 5,045 nuclear SNPs for the Medicinal Genomics StrainSEEK V1 dataset with use-type indicated below. Use-type of accessions include Type I=108, Type III=9, Type II=17 and Unknown=155 **(D)** Visualization of population structure and admixture from 20,566 nuclear SNPs for the Medicinal Genomics StrainSEEK V1 dataset using the fastSTRUCTURE software (k=2-5) with the optimal number of K being 3 using the silhouette method **(Fig. S9-10)**.

Phylogenetic trees were constructed in IQ-TREE (version 2.0.7 - Minh et al., 2020) with the -B 1000 flag for bootstrap support. Trees were visualized in FigTree (Version1.4.4 - http://tree.bio.ed.ac.uk/software/figtree/).

### Assembly of 126 whole chloroplast genomes

Chloroplast DNA was assembled using the Fast-Plast program, with default parameters - https://github.com/mrmckain/Fast-Plast. To explore haplo-group assignment a maximum likelihood phylogeny was constructed on 126 whole chloroplast sequences which were provided by LeafWorks Inc. Multiple sequence alignment was performed using MAFFT (version 7.475 - Katoh and Standley 2013) and using the ModelTest-NG software (Darriba et al. 2020) the GTR+G4 model was selected as the best substitution model. A phylogenetic tree was generated using IQ-TREE (version 2.0.7 - Minh et al., 2020) with the -B 1000 flag for bootstrap support. Trees were visualized in FigTree (Version1.4.4 - http://tree.bio.ed.ac.uk/software/figtree/).

## Results

### Commonalities across Datasets

*Cannabis* genetic diversity and population structure were explored using independent data sources, all of which were analyzed using the same pipeline (**Table 2**). All datasets were aligned to the same CBDRx reference genome (Grassa et al., 2021). Reanalysis allows for a cleaner comparison, as previous studies have used multiple reference genomes (e.g., Laverty et al., 2019; McKernan et al., 2020; van Bakel et al., 2011; Gao et al., 2020). There were not common SNPs across all datasets, when joint SNP calling was attempted. Therefore, each dataset was analyzed independently and each had a different number of SNPs (**Table 3-5)**. As sample sizes are robust, this suggests that the type of sequencing approach taken, library prep, sequencing depth, chromosomal coverage, and/or sample properties may bias the genetic diversity.

**Table 3.** SNP count per dataset pre and post filtering.

| Dataset | Sample (n) | Total # SNPs | # SNPs post filter (0.9) | # Bi-allelic SNPs | LD (0.2) | PC 1-6 (%) |
| --- | --- | --- | --- | --- | --- | --- |
| Phylos Biosciences | 845 | 1,620,202 | 385 | 383 | 292 | [1] 6.66 5.34 4.14 3.17 2.77 2.28 |
| Phylos Biosciences | 1,378 | 2,175,027 | 363 | 362 | 269 | [1] 8.14 4.61 4.18 3.41 2.91 2.76 |
| LeafWorks | 498 | 10,911,876 | 1,405 | 1400 | 520 | [1] 5.52 3.57 3.02 2.67 2.32 2.12 |
| Soorni *et al.* | 94 | 37,615,406 | 33,629 | 33,346 | 6,865 | [1] 4.84 2.85 2.12 1.96 1.83 1.68 |
| Sunrise Genetics | 25 | 7,502,178 | 6,329 | 6,284 | 1,604 | [1] 15.66 8.45 6.85 6.35 5.65 5.28 |
| Courtagen Life Sciences | 58 | 470,780,334 | 311 | 310 | 119 | [1] 6.16 4.83 4.59 4.47 4.26 3.86 |
| University of Colorado Boulder | 162 | 139,508,383 | 5,999 | 5,946 | 2,223 | [1] 5.40 3.64 3.12 2.57 2.45 2.24 |
| Medicinal Genomics 61 | 61 | 246,261,943 | 8,716 | 8,709 | 2,267 | [1] 4.95 3.89 2.75 2.53 2.39 2.32 |
| Medicinal Genomics StrainSEEK V1 | 289 | 121,471,853 | 20,566 | 20,454 | 5,045 | [1] 4.84 3.44 2.61 2.17 1.95 1.81 |

**Table 4.**
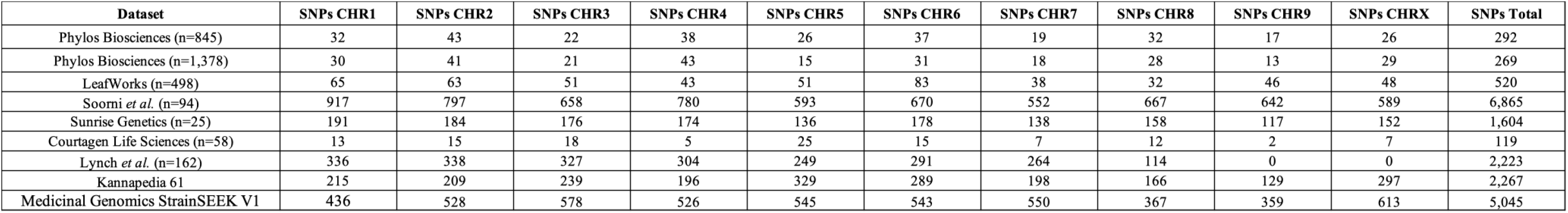
SNP counts for each dataset by chromosome following biallelic sorting and Linkage Disequilibrium prune at 0.2 and mapped to CBDRx (cs10) genome.

**Table 5.** Partition specific (Landrace and Domesticates) SNP count per dataset pre and post filtering.

|  | Accession # | Total # SNPs | # SNPs post filter | # Bi-allelic SNPs | LD (0.2) |
| --- | --- | --- | --- | --- | --- |
| LeafWorks (Landrace_101) | 101 | 4,761,034 | 2138 | 2131 | 1919 |
| LeafWorks (Domesticates_397) | 397 | 10,183,788 | 2096 | 2090 | 711 |
| LeafWorks (Hindu_Kush_6) | 6 | 835,656 | 4,304 | 4265 | 640 |
| LeafWorks (Lolab_Valley_4) | 4 | 502,884 | 853 | 850 | 170 |
| LeafWorks (Myanmar_Burma_4) | 4 | 617,666 | 2,204 | 2,186 | 384 |
| Phylos Biosciences (Landrace_107) | 107 | 1,027,670 | 267 | 266 | 219 |
| Phylos Biosciences (Domesticates_679) | 679 | 3,342,423 | 704 | 704 | 478 |

### Population Structure

Hierarchical Clustering of Principal Components (HCPC) identified three to seven clusters across datasets (**Fig. 1-2 and Fig. S1-6**). In the LeafWorks Inc. dataset, four groups were identified. When use-type was used to interpret the clusters there was some partitioning, with Group 1 being predominantly Hemp, but Groups 2/3 being largely type I (**Fig. 1A**). In the LeafWorks Inc. fastSTRUCTURE analysis there were large amounts of admixture regardless of use-type/market-class (**Fig. 1B**). Within the Phylos Biosciences datasets, hierarchical clustering identified five groups in the n=845 dataset (**Fig. 1C**) and three clusters in the n=1378 dataset (**Fig. S2B**). In the small Phylos Biosciences dataset (n=845) hierarchical clustering shows a concentrated number of Landrace (95 of 127), Hemp (14 of 17) and type I (11 of 48) samples in Group 1 (**Fig. 1C**). In Group 2 the majority of the type III samples are observed (30 of 48 – **Fig. 1C**). .fastSTRUCTURE analysis indicated less admixture in Landrace and Hemp samples as compared to type I samples (K=3/4/5), with some differentiation based on use-types observed (**Fig. 1D**). In the large dataset from Phylos Biosciences (n=1378 - **Fig. S2**), the hierarchical clustering shows a concentration of samples which have the designation of “Kush” in Group 1 (34 of 88 -**Fig. S2B**). The subsequent fastSTRUCTURE analysis for the Phylos Biosciences (n=1378) shows a similar pattern where Landrace samples show less admixture as compared to type I samples (K=4/5) (**Fig. S2C**). In the HCPC for the Phylos Biosciences dataset (n=1378) there is a concentration of samples with the designation “OG” (49/115 in Group 1 of HCPC) - **Fig. S2B**). In the Soorni et al dataset there was clear clustering by use-type in both analyses (**Fig. 2A-B**). The Medicinal Genomics StrainSEEK v1 dataset was partitioned into five groups (**Fig. 2C**). There was some clustering of specific genotypes (e.g. Blue Dream (n=11) in Group 1 of HCPC – **Fig. 2C**), but in fastSTRUCTURE analysis there were no clear trends in clustering observed across use-type (**Fig 2D**). For the Sunrise Genetics dataset (n=25) HCPC shows grouping of samples with the same names but no clear pattern in the fastSTRUCTURE clustering analysis (**Fig. S3C**). In the Lynch et al., 2016 dataset (n=162) there was some evidence of use-type but the pattern was not consistent (**Fig. S4B)**. For the Courtagen Life Sciences dataset (n=58), there was clustering by cultivar name (e.g. Kandy Kush (n=5) in Group 5 of HCPC) but not by use-type (**Fig. S5**). Within the Medicinal Genomics dataset (n=61) there was no clear clustering by use-type (**Fig. S6**). Across datasets there is no clear partitioning pattern based on use-type or based on accession name, this lack of pattern does not indicate a lack of population structure, but rather confirms the inconsistency in definitions of use-type and the fact that cultivar naming conventions do not reflect pedigrees.

### Phylogenetic Relationships

A Maximum Likelihood (ML) phylogeny was assembled for the new LeafWorks Inc. (n = 498) dataset which partitioned accessions into ten clades (**Fig. S7**). Clades 1 to 7 and clade 9 have bootstrap support of over 90, with clades 8 and 10 having low support (63 and 52, respectively). The majority of accessions (454 of 498) were in four clades (clades 4, 6, 9 and 10). There is not a clear pattern to which clade landrace samples are in (Clades 5=11 of 101; clade 9=24 of 101; clade10=52 of 101) with remaining individuals spread across the remaining clades - **Fig. S7)**. There did not appear to be clear use-type partitioning in the phylogeny. Within the chloroplast data, two clades were identified, clade 1 (n= 2) and clade 2 (n=124) (**Fig. S8**). However, with low support for the majority of samples it is possible that additional groupings within this might be possible.

### Exploration of nucleotide diversity and geographic partitioning Landraces

The LeafWorks Inc. (n=498) and Phylos Biosciences (n=845) datasets both contained known landrace and modern cultivars (**Table 1; Table S2-S3**). The LeafWorks Inc. dataset contained 101 landrace samples and 397 known modern accessions (**Table S1**) and the Phylos Biosciences dataset contained 127 landrace samples and 718 modern accessions (**Table S2**). Clustering patterns were similar in the two datasets (**Fig. 3**). Within both datasets nucleotide diversity differences were explored between landrace and domesticated samples using a 10 kb sliding window (**Fig. 4**), revealing many genomic regions that differed between modern cultivars and landraces. A subset of landraces in the LeafWorks Inc. dataset contained geo-references, allowing for an exploration of structure based on geography. There was geographic clustering with the Lolab Valley and Hindu Kush samples (**Fig. 5C**). There were low levels of admixture based on the three geographic regions (**Fig. 5E**), indicating that despite being geographically close populations remained isolated. Differences in nucleotide diversity in these geographically distinct populations were observed on chromosomes 5, 6, and 7 (**Fig. 5D)**, and these genomic regions may hold locally adaptive genes and may be useful sources of variation for breeding.

**Fig. 3.**
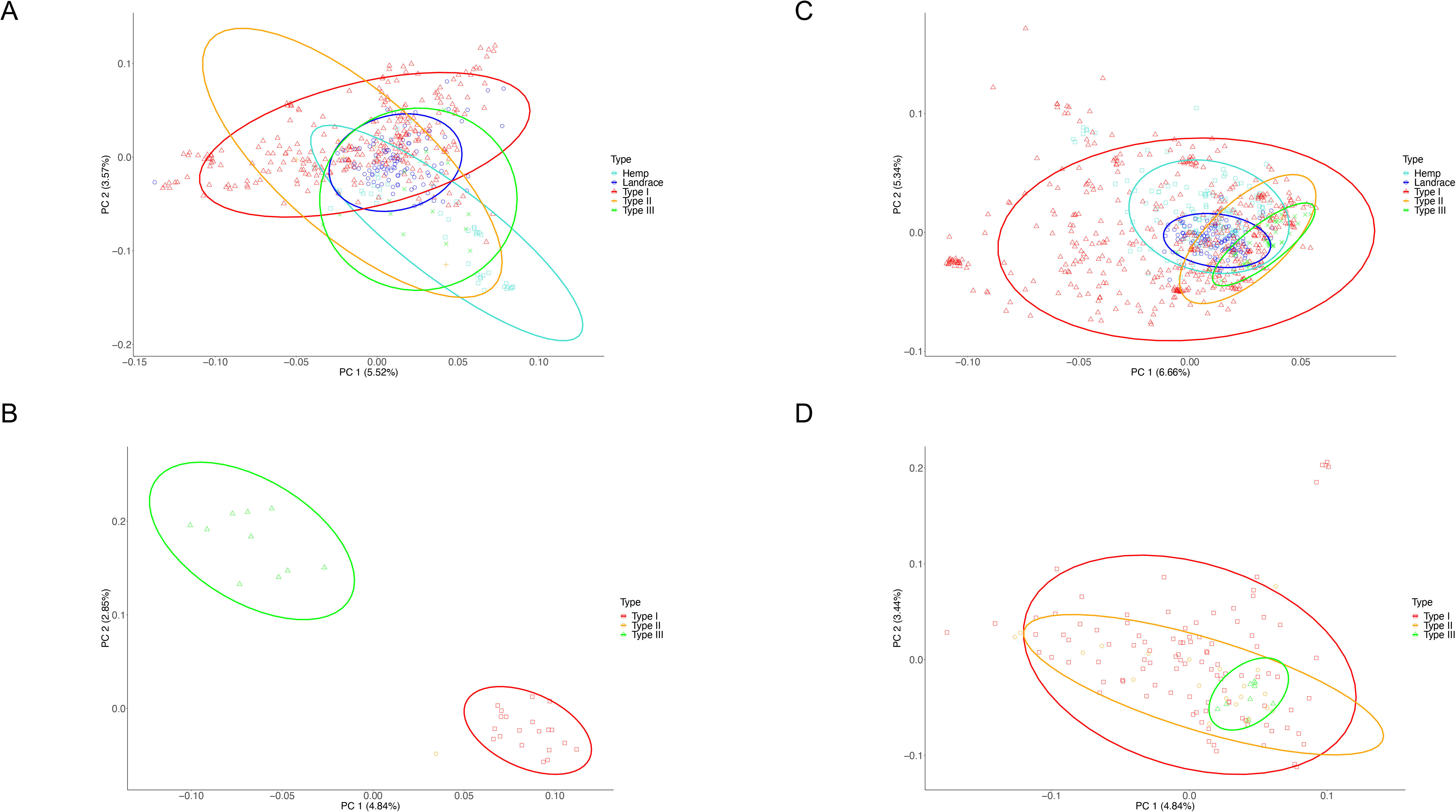
Examination of use-type association across datasets **(A)** Principal component analysis (PCA) from 520 nuclear SNPs for the LeafWorks Inc. dataset **(B)** PCA from 213 SNPs Phylos Biosciences(n=845) dataset **(C)** PCA from 6,865 nuclear SNPs for the Soorni et al. 2017 dataset where cannabinoid content could be determined due to recent publication for 31/94 samples. (**D)** PCA from 5,045 nuclear SNPs for the Medicinal Genomics StrainSEEK V1 dataset.

**Fig. 4.**
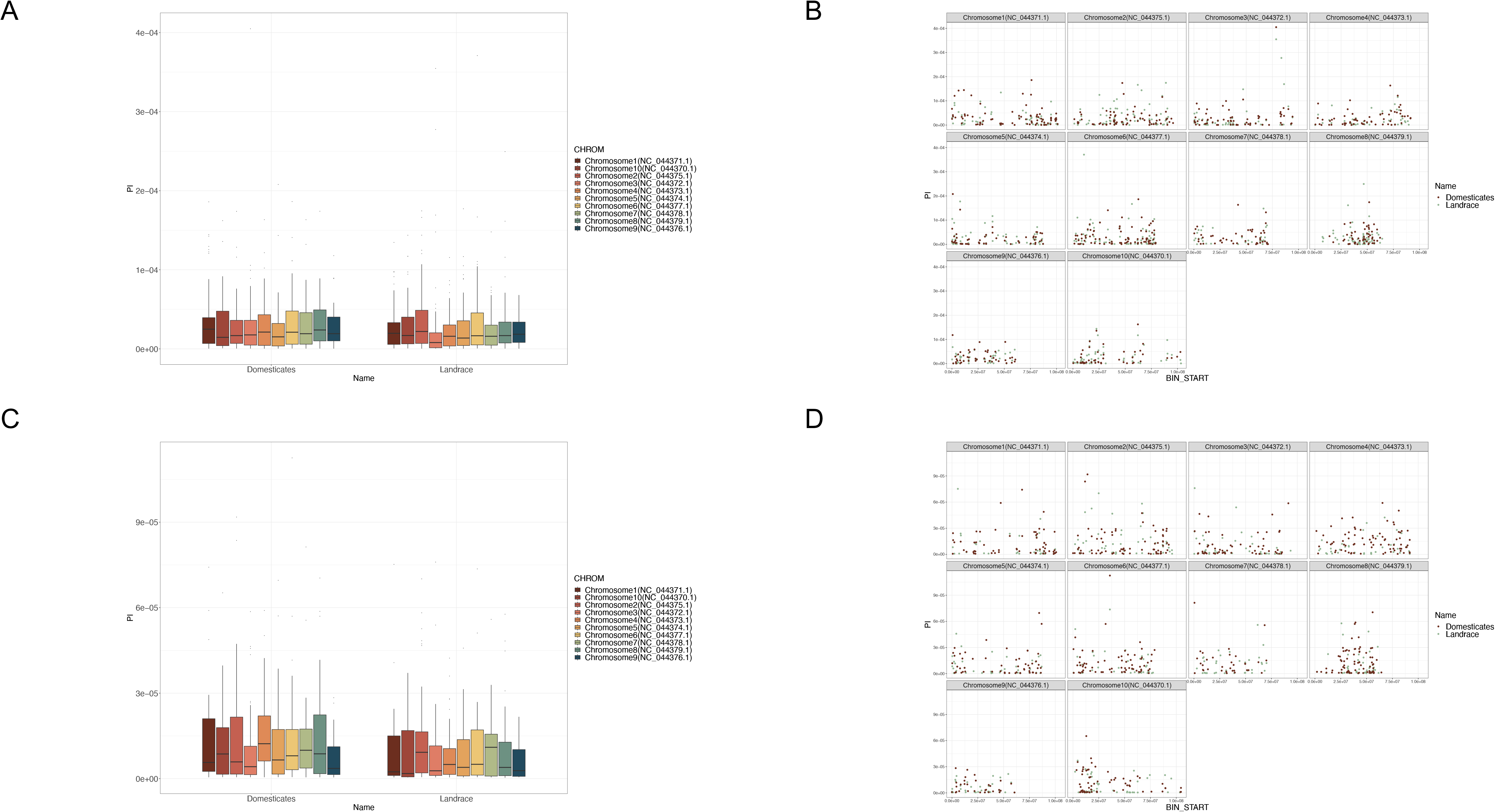
Nucleotide diversity as examined by a 10kb sliding window for landrace and domesticated partitions for the LeafWorks Inc. and Phylos Biosciences datasets **(A)** Nucleotide diversity by chromosome and **(B)** across chromosome length for Domesticated (n=397, 2,096 SNPs) and Landrace (n=101, 2,131 SNPs) samples for the LeafWorks Inc. dataset **(C)** Nucleotide diversity by chromosome and (D) across chromosome length for Domesticated (n=718, 749 SNPs) and Landrace (n=127, 566 SNPs) samples for the Phylos Biosciences dataset.

**Fig. 5.**
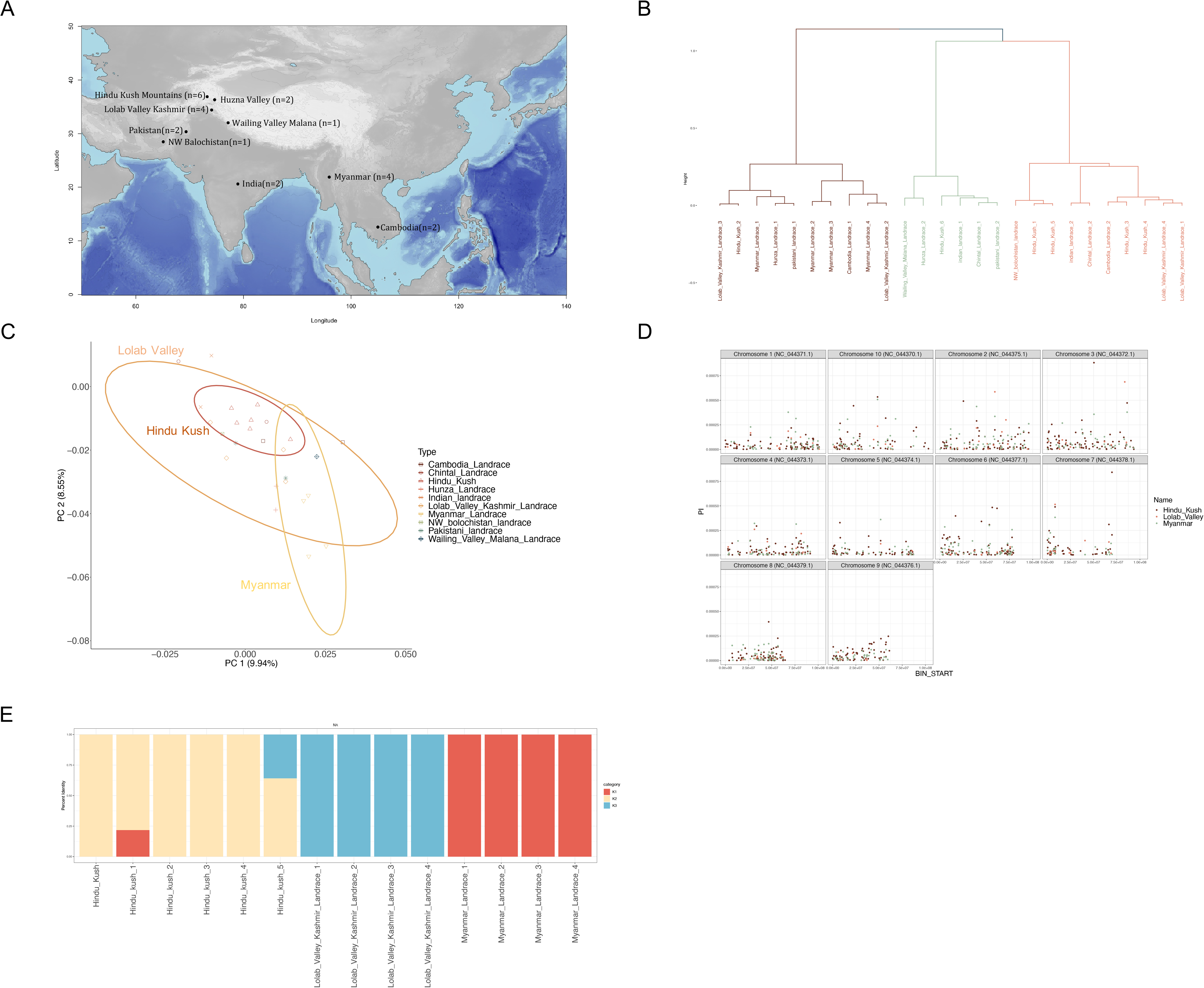
Landrace accessions from the LeafWorks Inc. dataset show separation between Indian and Myanmar populations **(A)** Map detailing the locations of landrace accessions, highlighted are the Hindu Kush Mountains, Lolab Valley and Myanmar **(B)** Hierarchical cluster dendrogram based on 304 SNPs (LD 0.2) across 26 samples of known and trusted origin (**C)** PCA based on 304 SNPs with geographical locations of samples as indicated **(D)** Nucleotide diversity comparison between Hindu Kush Mountains (n=6, 4,304 SNPs), Lolab Valley (n=4, 853 SNPs) and Myanmar (n=4, 2,204 SNPs) as examined by a 10kb sliding window **(E)** Visualization of population structure and admixture using the fastSTRUCTURE software (k=3) with the optimal number of K being 3 using the silhouette method.

### Core collection identification

Plant breeding relies on the available genetic variation within a given germplasm collection or breeding program. A core collection is a representative subset of a germplasm collection which attempts to capture the majority of the genetic diversity in that collection (Frankel 1984). Using genetic distance, the 25 most diverse samples were selected from each dataset (with the exception of the Sunrise dataset where 10 samples were selected). These samples represent a core collection for each specific dataset (**Table S10**).

## Discussion

Species descriptions in the *Cannabis* genus have been based on morphology and chemistry (Clarke & Merlin, 2013; Onofri & Mandolino, 2017; Lewis et al., 2018; McPartland, 2018; Garfinkel et al., 2021; Smith et al., 2022). While three putative species have been described in the *Cannabis* genus, genetic studies have not supported these delineations, instead observing a monotypic genus (Clarke & Merlin, 2013; Sawler et al., 2015; Lynch et al., 2016; Schwabe & McGlaughlin, 2019). Much effort has been made exploring use-type/marketing class as markers of population stratification (Clarke & Merlin, 2013; Small, 2015). Here we continue this tradition, by exploring population structure in nine different collections of *Cannabis* that consisted of privately bred THC-dominant, public hemp samples and landrace accessions. Understanding genetic diversity within each individual collection aides in understanding population history and helps in developing strategies for future breeding. In particular, establishing the number of distinct populations may help reduce the number of individuals that need to be tested for the development of hybrid cultivars (Carlson et al., 2021).

### Understanding population structure

Previous work has used various reference genomes (Lynch et al., 2016; Soorni et al., 2017; Laverty et al., 2019; Jin et al., 2021) and this reflects the current predicament within the industry where standards are still in development. Here a single reference genome (CBDRx - Grassa et al. 2021) was used to facilitate comparison; however, it is acknowledged that this can create reference bias impacting the examination of some questions. Reference limitations are being addressed through the utilization of pangenomes and are increasingly becoming available for many crop species (Hübner et al., 2019; Li et al., 2021; Della Coletta et al., 2021). Future work to develop a *Cannabis* pangenome would be of great utility to the community.

*Cannabis* has historically used morphological and ethnographic data to delineate populations not genetic data. Creating genetic profiles to cluster accessions and conduct phylogenetic analysis facilitates using classic use-type to understand accession relationships (**Fig 3 and Fig. S7**) and offers a perspective on how *Cannabis* populations may have been influenced by human mediated selection for important traits (**Fig. S2-6**). The LeafWorks Inc. and Soorni et al., 2017 datasets exhibited more use-type separation (**Fig. 3A/3C**) than other datasets. A broader distribution of genetic variation in Type I cultivars was observed in multiple datasets (**Fig. 3A-B**). This may be indicative of the purported large-scale hybridization that is thought to have occurred in Type I cultivars in the United States after 1960 (Clarke & Merlin, 2016). Alternatively, this higher genetic diversity in Type I cultivars could represent convergent selection, with each lineage being bred in isolation and now released back to the market as regulations relax. However, the lack of available pedigree records makes it difficult to reconcile these two alternative hypotheses. The other datasets did not show clear relationships with use-type.

While the *Cannabis* genus has been described with the presence of one, two, three, and even up to seven proposed species and subspecies (Linnaeus, 1753; Lamarck, 1785; Vavilov & Bukinich, 1929; Schultes et al., 1974; Small & Cronquist, 1976; Hillig, 2004; Clarke & Merlin, 2013; McPartland & Guy 2014), contemporary genetic studies have not supported these polytypic classifications delineations which are primarily rooted in morphological and geographical data. While modern genetic studies consistently do not support previously suggested species delineation (Gilmore et al., 2007; Sawler et al., 2015; Lynch et al., 2016; Small, 2015; Zhang et al., 2018; Schwabe & McGlaughlin, 2019; McPartland & Small 2019; Roman et al., 2019; Henry et al. 2020; Ren et al 2021; Schwabe et al., 2021; Vergara et al. 2021; Woods et al., 2023), they do support multiple potential populations within the genus. Despite these findings, the prevailing taxonomic treatment of the *Cannabis* genus tends to favor a monotypic classification. Several publications have proposed delineations in the relationship between use-type and population structure (Gilmore et al., 2007; Roman et al., 2019; Zhang et al., 2018; Henry et al., 2020; Ren et al., 2021; Woods et al., 2023). This suggests that collection origin and accurate passport data greatly impact the population structure observed. High levels of hybridization and shared ancestry may all contribute to the relatively shallow population structure observed in some datasets (**Fig. 1D**) and hamper the ability to clearly differentiate populations.

Landrace samples are distributed throughout population clusters and across the phylogenies, however the number of landrace samples in a particular partition appear to be affected by the germplasm sampling (**Fig. 1 and Fig. S7**). When landraces were analyzed with modern cultivars, they were broadly distributed across clades suggesting that modern cultivars in the same clade share the most ancestry with the landraces in the same clade (**Fig. 1A-B**). Landraces did not cluster with a particular use-type. In a subset of georeferenced samples (n=26) hierarchical clustering revealed three geographically discrete landrace populations appear to be quite distinct from one another with minimal admixture (**Fig. 5B/E**). The subset landraces with georeferences and which were geographically separated, clustered distinctly when analyzed separately (**Fig. 5E**). While landraces were defined based on metadata and a history of being grown in a specific geography, limitations on passport data cloud inference. Without new collections and legitimate chain of custody documentation, this likely cannot be addressed.

### Inconsistent Naming

The naming problem in *Cannabis* refers to the unreliable naming of cultivars which frequently do not reflect accession pedigree causing problems for both the producer and consumer. Name fidelity was explored using the twelve ‘Blue Dream’ samples in the LeafWorks Inc. dataset (**Fig. S7**). Of these, 7/12 placed in clade 1 (blue_dream samples #1, 3, 4, 6, 7, 9 & 10), 1 sample in clade 6 (blue_dream_5), clade 9 (blue_dream_11) and clade 10 (blue_dream_2). The remaining two samples (blue_dream # 8 & 12) were unplaced in the phylogenetic tree. Cultivars that show consistent placement within a phylogeny have a high likelihood of name accuracy. This exemplifies the naming problem, where only 58% of the samples appeared to be similar. This data further supports previous work demonstrating misconceptions in strain reliability and which further showed that the marketing varieties of *Cannabis* as “indica” and “sativa” does not appear to have genetic support (Schwabe & McGlaughlin, 2019). Further work is needed to determine how pervasive the naming problem is. This work also highlights the importance of genetics to inform label claims, which will be particularly crucial in the event of legalization when the Federal Drug Administration would require accurate plant label claims as it does for all other natural products sold in the United States. As sequencing costs continue to decrease, genomic approaches for understanding *Cannabis* naming will likely become standard practice and could overcome the challenge of clone and cultivar misidentification.

### Strategic use of Germplasm for Breeding

Variation in cannabinoid content is genetically complex and potentially affected by the environment (Lydon et al., 1987; Campbell et al., 2020; Caplan et al., 2019; Toth et al., 2021). Breeding with a focus on a particular use-type could help to ensure consistency in secondary chemistry and incorporating an assessment of admixture or hybridization in this selection may expedite the time taken to reach population stability. Coupling plant phylogeny with metabolomics could facilitate the identification of plants with unique genetic and secondary chemistries and would provide unique market classes (Stone et al. 2020).

Breeding targets in the future will likely focus on the common traits of disease and pest resistance but will also likely need to maintain certain metabolite content to ensure use-type. There is potential value in exploring if specific SNP markers can be identified to differentiate use-type as this can inform parental choice in plant breeding programs. Expanding the use of genome wide markers will not only help to characterize populations but can also help establish preliminary partitioning of samples into potential heterotic groups. Population stratification and use-type categorization have already found applications in hybrid breeding efforts (Carlson et al., 2021). To establish new patterns of heterosis, it has been proposed that a practical starting point could involve segregating individuals based on genetic distance, with a threshold set at >0.4 (Govindaraju 2019). This approach can be empirically tested within any germplasm collection, whether it’s publicly or privately held, to identify effective patterns for achieving improved breeding outcomes.

Another option will be to use evolutionary plant breeding (EPB) to help maintain diversity and stability of a crop in a specific environment leveraging natural selection (Merrick et al. 2020). This has been used to aid in hybridization (Dreiseitl, 2020). Here landrace sampling was limited (**Tables S1-9**), but a more thorough characterization of *Cannabis* landrace populations would facilitate use of this approach. The history of prohibition and local cultivar development suggests that there is a large possibility of biopiracy (e.g. unauthorized exploitation or theft of valuable genetic resources or traditional knowledge) with respect to the developing industry. It will be important to develop equitable distribution and ensure that local communities benefit from the work their communities have done in the past and be in compliance with The International Treaty on Plant Genetic Resources for Food and Agriculture (Cooper, 2002).

### Public Data Implications

The relaxing of governmental regulations and decrease in sequencing costs technologies have made it possible to genotype many different germplasm collections over the last decade (**Table 2**). Public-private partnerships offer a route to harness the diverse resources and expertise present in both sectors and provide a useful mechanism to advance *Cannabis* science (Ferroni & Castle, 2011). Ensuring data standards are upheld and that metadata are available will make the increasing amount of data available useful to many different researchers (Chao, 2014). The ability to analyze the data requires accurate metadata and while this problem is not unique to *Cannabis*, it is acutely problematic in any species that has high economic value and limited foundational genomic resources. When working with public data sources care must be taken in the cross comparison of specific datasets as the amount of shared germplasm and data quality can influence the breadth and inference potential of the analysis (Williamson et al. 2021). Additionally, when expanding these observations to conclusions about the genus as whole, it is important to carefully consider germplasm sampling bias which limits the direct comparison, which may result in limited or no shared SNPs across datasets (Zimmerman et al., 2020). It is evident that the selection of sequencing technology, such as short-read amplicon sequencing or genome-wide sequencing, can substantially alter the capacity for making inferences and can have a notable impact on the value of some genomic statistics (Evangelou and Ioannidis, 2013; Marchi et al., 2021). In this analysis it is very likely that due to the high numbers of potential Type I plants, sampling of male *Cannabis* plants has been largely unobserved. This is because Type I plants typically consist of female flowers exclusively, with males often being removed from cultivation.

### Future perspectives

Nine datasets were explored to understand population structure in *Cannabis*, identifying inconsistent genetic clustering with use-type. The inconsistency of use-type as a predictor of relatedness implies that it may be a collection specific association, or that the relatively simple inheritance of tetrahydrocannabinolic acid synthase (THCAS) may obfuscate background genetic relationships. With the legal status of *Cannabis* now shifting, researchers can begin to examine the effects of prohibition on extant *Cannabis* varieties and keep better records while developing new cultivars. In the United States, prohibition may have created closed gene pools through the breeding of limited germplasm facilitated by limiting plant exchange. Limited genetic diversity in breeding may have had a role to play in the increased potency of *Cannabis* varieties over time, with increases in THC content from ∼4% in 1995 to ∼12% in 2014 reported (El Sohly et al. 2016). Analogous to this in the wild, repeated range contractions during the Holocene are thought to have resulted in repeated genetic bottlenecks and likely initiated incomplete allopatric speciation which has led to differences between European (CBD-dominant - Type III) and Asian (THC-Dominant - Type I) *Cannabis* populations (McPartland 2018).

The study of *Cannabis* genetics and its population structure is influenced by historical factors like prohibition and contemporary breeding practices. Genome sequencing technologies play a pivotal role in shaping our understanding of *Cannabis* genetic diversity. The debate over species and subspecies classifications persists, with genetic research consistently challenging traditional delineations. Importantly, the identification of population stratification and genetic markers holds promise for enhancing breeding efforts, particularly in developing new heterotic patterns. Additionally, while the industry burgeons, concerns regarding biopiracy and the preservation of genetic diversity remain salient. Despite the challenges posed by inconsistent naming conventions and limited sampling, ongoing research efforts continue to shed light on the intricate genetic landscape of *Cannabis,* with significant implications for its future cultivation, medicinal use, and industrial applications.

## Acknowledgements

We would like to thank Mr. Robert Connell Clarke for his curation of the use-type associations for the Phylos Biosciences (n=1378) dataset as well as for valuable discussions and insights.

## Availability of data and material

data are available upon reasonable request to the corresponding author.

## Code availability

code are available at https://github.com/ahmccormick and at https://figshare.com/authors/Anna_H_McCormick/17741367

## Author contributions

*Conceptualization:* AHMC, KH, MBK, NB, RRM, KL, EJK, *Formal Analysis:* AHMC, RRM, *Figure Preparation:* AHMC, *Manuscript Drafting:* AHMC, *Writing and Reviewing Manuscript:* AHMC, KH, MBK, NB, RRM, KL, EJK.

## Supplemental Figure Legends

**Fig. S1** Dataset overview **(A)** Nucleotide diversity examined by a 10kb sliding window for all 9 genomic datasets for *Cannabis sativa* L. **(B)** Nucleotide diversity across the length of the 10 chromosomes for all 9 genomic datasets.

**Fig. S2** Nuclear SNP analysis for the Phylos Biosciences (n=1,378) dataset. Clustering was conducted based on nuclear genetic SNPs while reported use-type within the dataset is below in solid bars to facilitate interpretation based upon community standards **(A)** PCA by use-type based on 269 nuclear SNPs. Use-type associations include THC-Dominant (Type I) (n=996) CBD-Dominant (Type III) (n=87), Hemp (n=215), Landrace (n=78) and Unknown (n=2) **(B)** Hierarchical cluster dendrogram with use-type indicated below (C) Visualization of population structure and admixture using the fastSTRUCTURE software (k=2-5) with the optimal number of K being 3 using the silhouette method.

**Fig. S3** Nuclear SNP analysis for the Sunrise Genetics dataset for 25 samples. Clustering was conducted based on nuclear genetic SNPs while reported use-type within the dataset is below in solid bars to facilitate interpretation based upon community standards **(A)** PCA by use-type based on 1,604 nuclear SNPs. Use-type associations include THC-Dominant (Type I) (n=38) and Unknown (n=12) **(B)** Hierarchical cluster dendrogram with use-type indicated below **(C)** Visualization of population structure and admixture using the fastSTRUCTURE software (k=2-5) with the optimal number of K being 3 using the silhouette method.

**Fig. S4** Nuclear SNP analysis for the Lynch et al., 2016 dataset for 162 samples. Clustering was conducted based on nuclear genetic SNPs while reported use-type within the dataset is below in solid bars to facilitate interpretation based upon community standards **(A)** PCA by use-type for 162 samples from 2,223 SNPs. Type associations include Hemp (n=1), Landrace (n=1), THC-Dominant (Type I) (n=162), CBD-Dominant (Type III) (n=11), THC:CBD (Type II) (n=2) and Unknown (n=21) **(B)** Hierarchical cluster dendrogram with use-type indicated below **(C)** Visualization of population structure and admixture using the fastSTRUCTURE software (k=2-5) with the optimal number of K being 2 using the silhouette method.

**Fig. S5** Nuclear SNP analysis for the Courtagen Life Sciences dataset for 58 samples. Clustering was conducted based on nuclear genetic SNPs while reported use-type within the dataset is below in solid bars to facilitate interpretation based upon community standards **(A)** PCA by use-type based on 119 nuclear SNPs. Use-type associations include Hemp (n=1), THC-Dominant (Type I) (n=41), CBD-Dominant (Type III) (n=11) and Unknown (n=5) **(B)** Hierarchical cluster dendrogram with use-type indicated below **(C)** Visualization of population structure and admixture using the fastSTRUCTURE software (k=2-5).

**Fig. S6** Nuclear SNP analysis for the Medicinal Genomics 61 dataset for 61 samples. Clustering was conducted based on nuclear genetic SNPs while reported use-type within the dataset is below in solid bars to facilitate interpretation based upon community standards **(A)** PCA by use-type based on 2,267 nuclear SNPs. Use-type associations include Hemp (n=1), THC-Dominant (Type I) (n=47), CBD-Dominant (Type III) (n=5) and Unknown (n=9) (**B)** Hierarchical cluster dendrogram with use-type indicated below **(C)** Visualization of population structure and admixture using the fastSTRUCTURE software (k=2-5) with the optimal number of K being 3 using the silhouette method.

**Fig. S7** Maximum Likelihood tree for the LeafWorks Inc. dataset constructed from 1,405 nuclear SNPs from 498 samples. Modeltest-ng revealed the TIM2+G4 as the best fit substitution model and IQ-Tree software was used for phylogenetic inference. Blue Dream samples (n=12) are highlighted in blue at the branch tips. Use-type for individual samples is additionally indicated.

**Fig. S8** Maximum Likelihood phylogenetic tree for 126 whole chloroplast assemblies. Individuals were aligned using MAFFT. Modeltest-NG revealed the GTR+G4 as the best fit substitution model and IQ-Tree software was used for phylogenetic inference. The resultant tree was visualized using FigTree (Version 1.4.4).

**Figure S9** Examining optimal K number across the datasets using the Elbow Method **(A)** LeafWorks Inc. dataset **(B)** Phylos Biosciences dataset (n=845) **(C)** Soorni dataset (n=94) **(D)** Medicinal Genomics StrainSEEK V1 (n=289) **(E)** Phylos Biosciences dataset (n=1378) **(F)** Sunrise Genetics (n=25) **(G)** Colorado dataset (n=162) **(H)** Courtagen dataset (n=58) **(I)** Kannapedia 61 dataset (n=61) **(J)** LeafWorks Inc. landrace samples (n=14).

**Figure S10** Examining optimal K number across the datasets using the Silhouette Method **(A)** LeafWorks Inc. dataset **(B)** Phylos Biosciences dataset (n=845) **(C)** Soorni dataset (n=94) **(D)** Medicinal Genomics StrainSEEK V1 (n=289) **(E)** Phylos Biosciences dataset (n=1378) **(F)** Sunrise Genetics (n=25) **(G)** Colorado dataset (n=162) **(H)** Courtagen dataset (n=58) **(I)** Kannapedia 61 dataset (n=61) **(J)** LeafWorks Inc. landrace samples (n=14).

## Table Legends

**Table S1.** Cultivar name, use-type, clade association and domestication classifications for the LeafWorks Inc. data set.

**S2.** SSR ID, Cultivar name, use-type and domestication classifications for the Phylos Biosciences (n=845) data set.

**Table S3.** SSR ID, Cultivar name, use-type, clade association and domestication classifications for the Phylos Biosciences (n=1,378) data set.

**Table S4.** SSR ID, Cultivar name, Chemistry Type and and HCPC group for the Soorni et al. 2017 data set.

**Table S5.** Sample ID, RSP ID, Cultivar name and use-type association for Medicinal Genomics (n=753) data set.

**Table S6.** SSR ID, Cultivar name and use-type association for the Sunrise Genetics data set.

**Table S7.** SSR ID, Cultivar name and use-type association for Lynch et al., 2016 data set.

**Table S8. S**SR ID, Cultivar name and use-type association for the Courtagen Life Sciences data set.

**Table S9.** Sample ID, Cultivar name and use-type association for Medicinal Genomics (n=61) data set.

**Table S10.** Core collections for the nine datasets.

