## Supplementary material for "Examining population structure across multiple collections of Cannabis": Supplemental_Figures_1-25-24.pdf

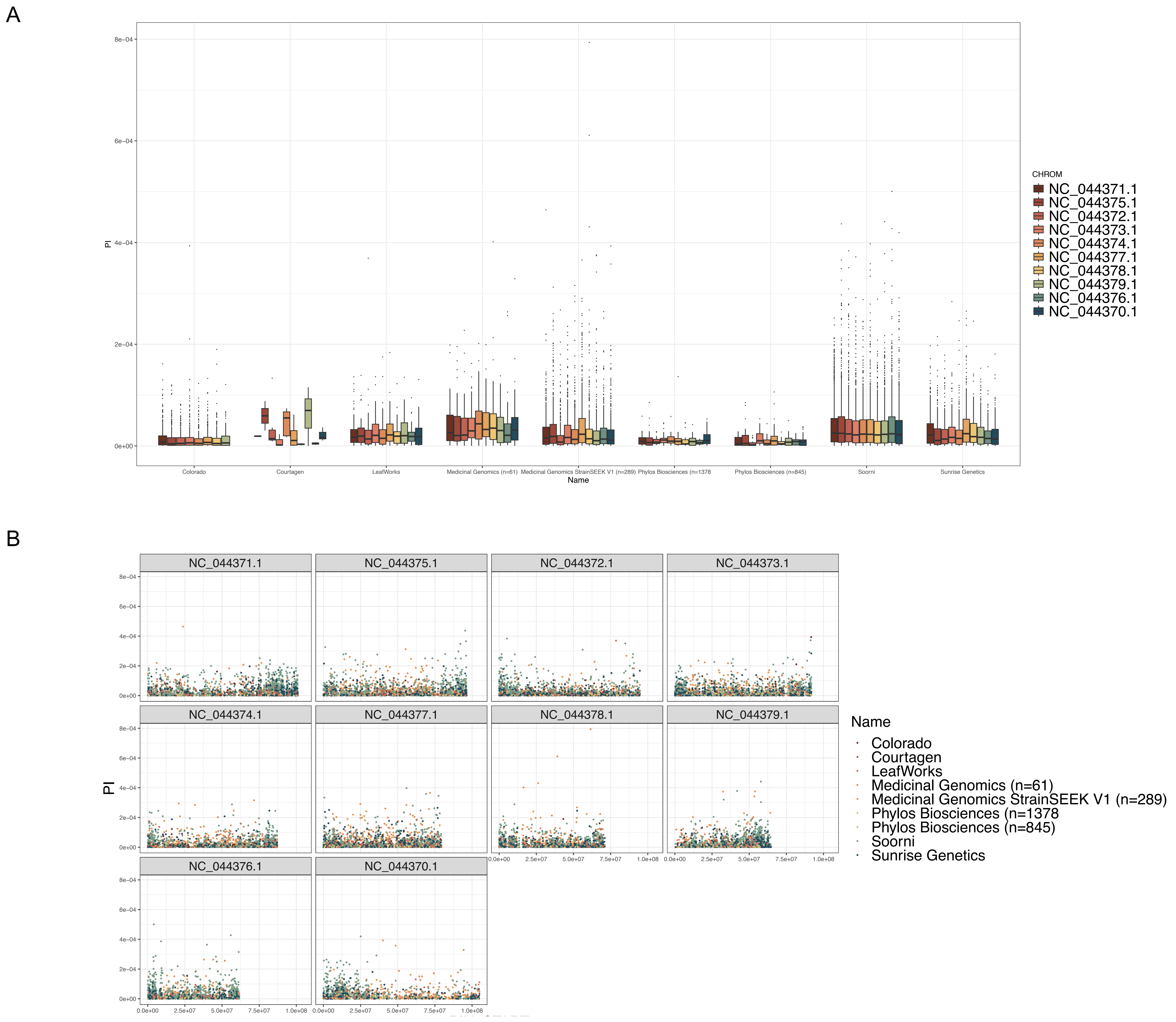


**Supplemental Figure 1** Dataset overview **(A)** Nucleotide diversity examined by a 10kb sliding window for all 9 genomic datasets for *Cannabis sativa* L. **(B)** Nucleotide diversity across the length of the 10 chromosomes for all 9 genomic datasets.

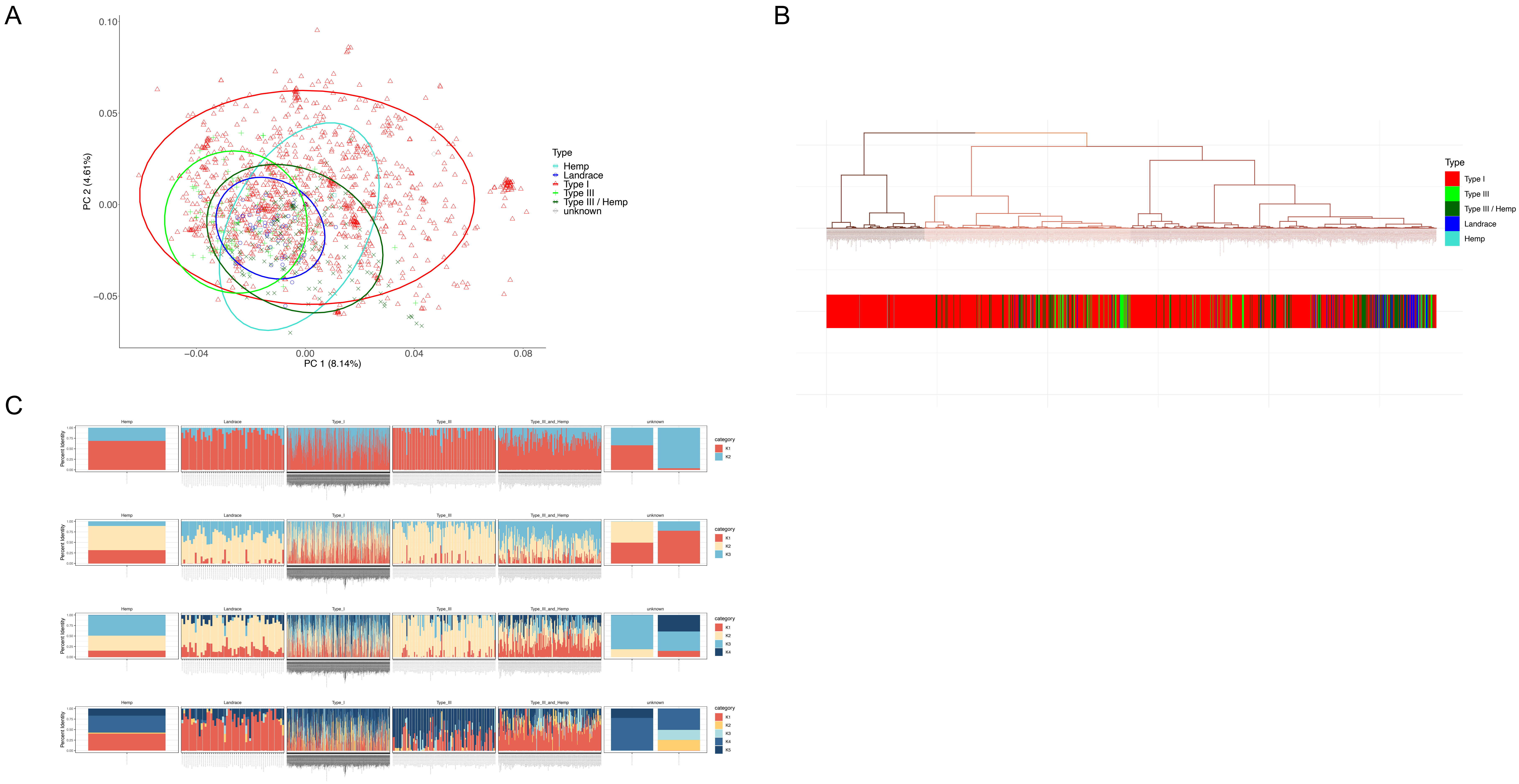


**Supplemental Figure 2** Nuclear SNP analysis for the Phylos Biosciences (n=1,378) dataset. Clustering was conducted based on nuclear genetic SNPs while reported use-type within the dataset is below in solid bars to facilitate interpretation based upon community standards **(A)** PCA by use-type based on 269 nuclear SNPs. Use-type associations include THC-Dominant (Type I) (n=996) CBD-Dominant (Type III) (n=87), Hemp (n=215), Landrace (n=78) and Unknown (n=2) **(B)** Hierarchical cluster dendrogram with use-type indicated below (C) Visualization of population structure and admixture using the fastSTRUCTURE software (k=2-5) with the optimal number of K being 3 using the silhouette method.

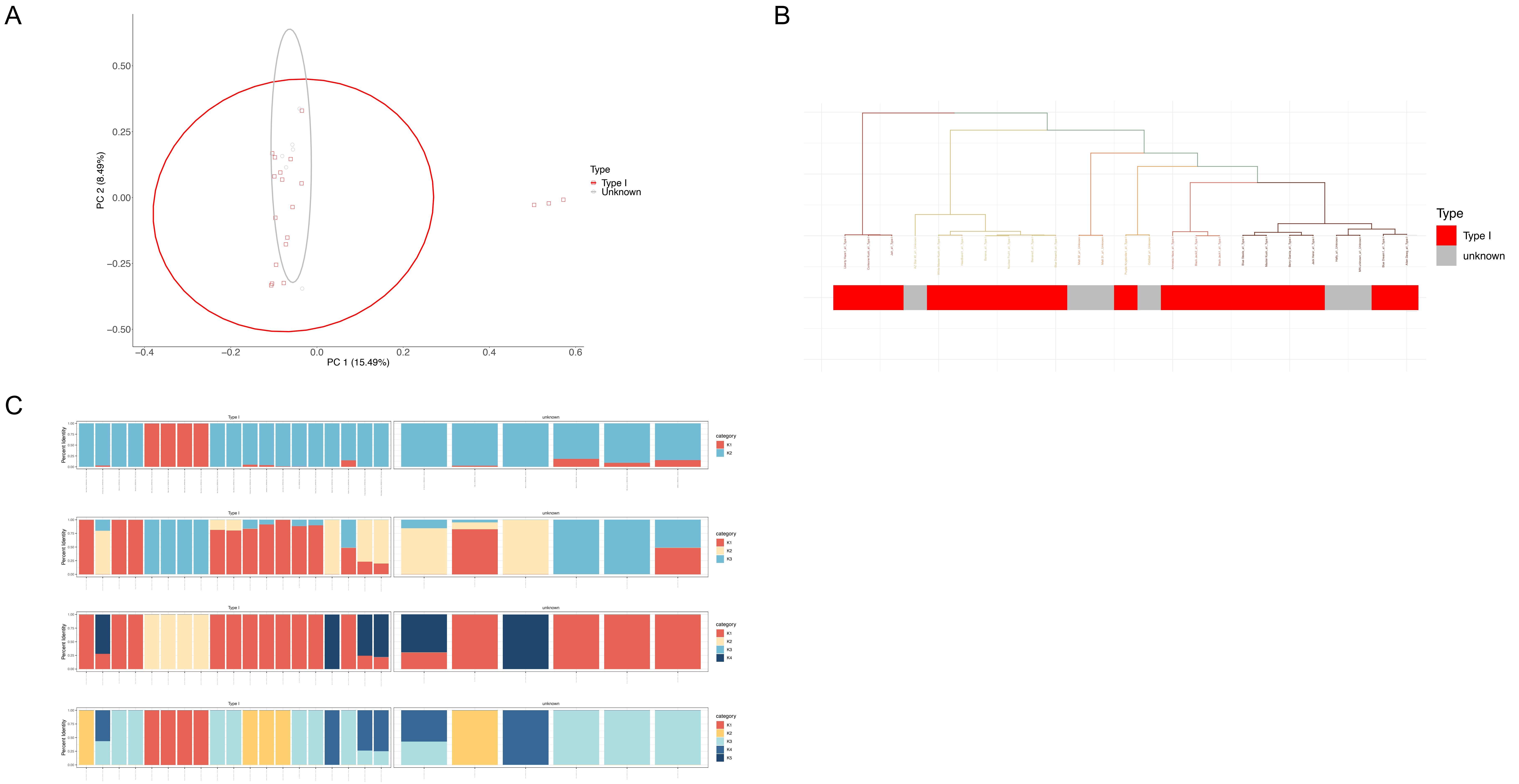


**Supplemental Figure 3** Nuclear SNP analysis for the Sunrise Genetics dataset for 25 samples. Clustering was conducted based on nuclear genetic SNPs while reported use-type within the dataset is below in solid bars to facilitate interpretation based upon community standards **(A)** PCA by use-type based on 1,604 nuclear SNPs. Use-type associations include THC-Dominant (Type I) (n=38) and Unknown (n=12) **(B)** Hierarchical cluster dendrogram with use-type indicated below **(C)** Visualization of population structure and admixture using the fastSTRUCTURE software (k=2-5) with the optimal number of K being 3 using the silhouette method.

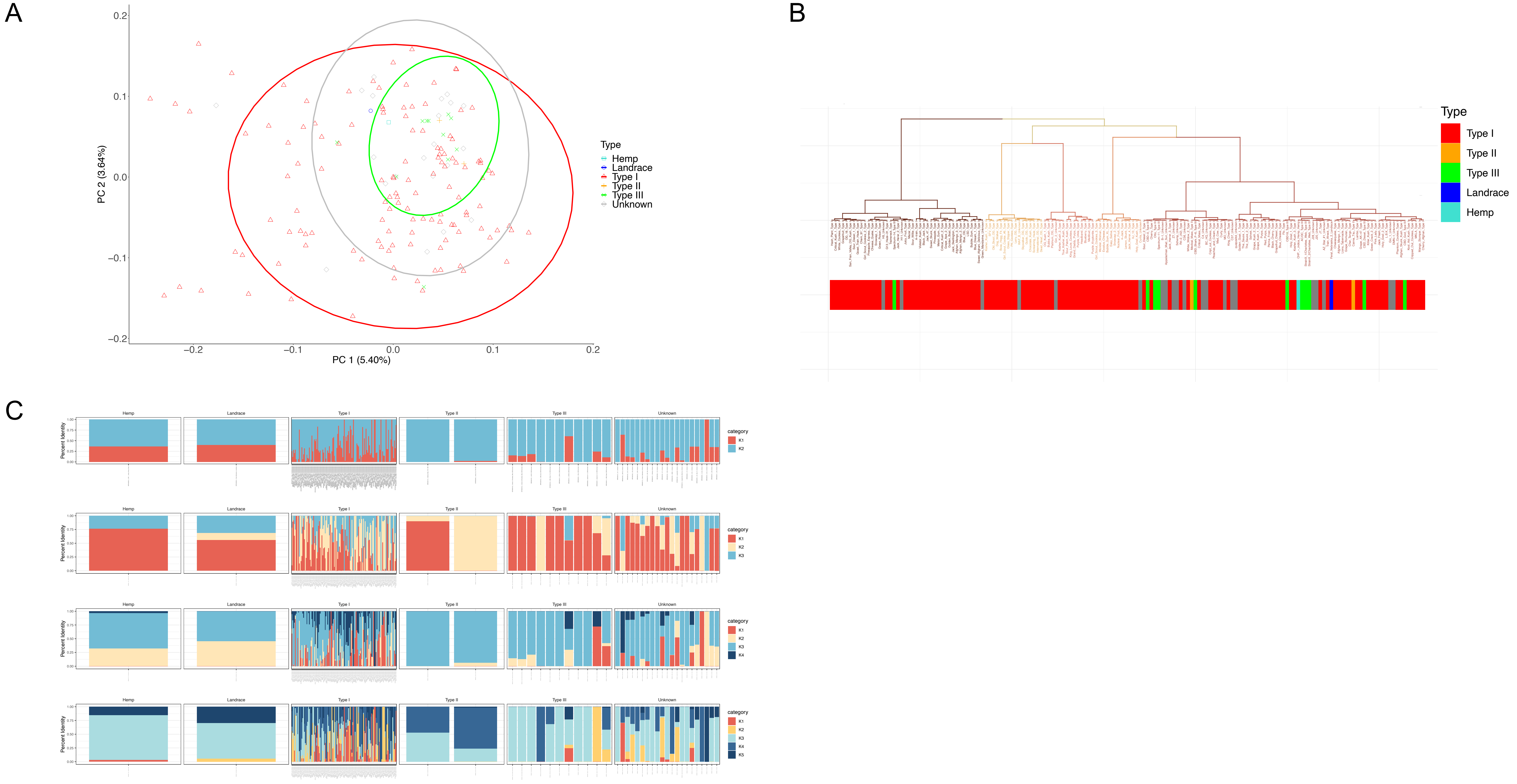


**Supplemental Figure 4** Nuclear SNP analysis for the Lynch et al., 2016 dataset for 162 samples. Clustering was conducted based on nuclear genetic SNPs while reported use-type within the dataset is below in solid bars to facilitate interpretation based upon community standards **(A)** PCA by use-type for 162 samples from 2,223 SNPs. Type associations include Hemp (n=1), Landrace (n=1), THC-Dominant (Type I) (n=162), CBD-Dominant (Type III) (n=11), THC:CBD (Type II) (n=2) and Unknown (n=21) **(B)** Hierarchical cluster dendrogram with use-type indicated below **(C)** Visualization of population structure and admixture using the fastSTRUCTURE software (k=2-5) with the optimal number of K being 2 using the silhouette method.

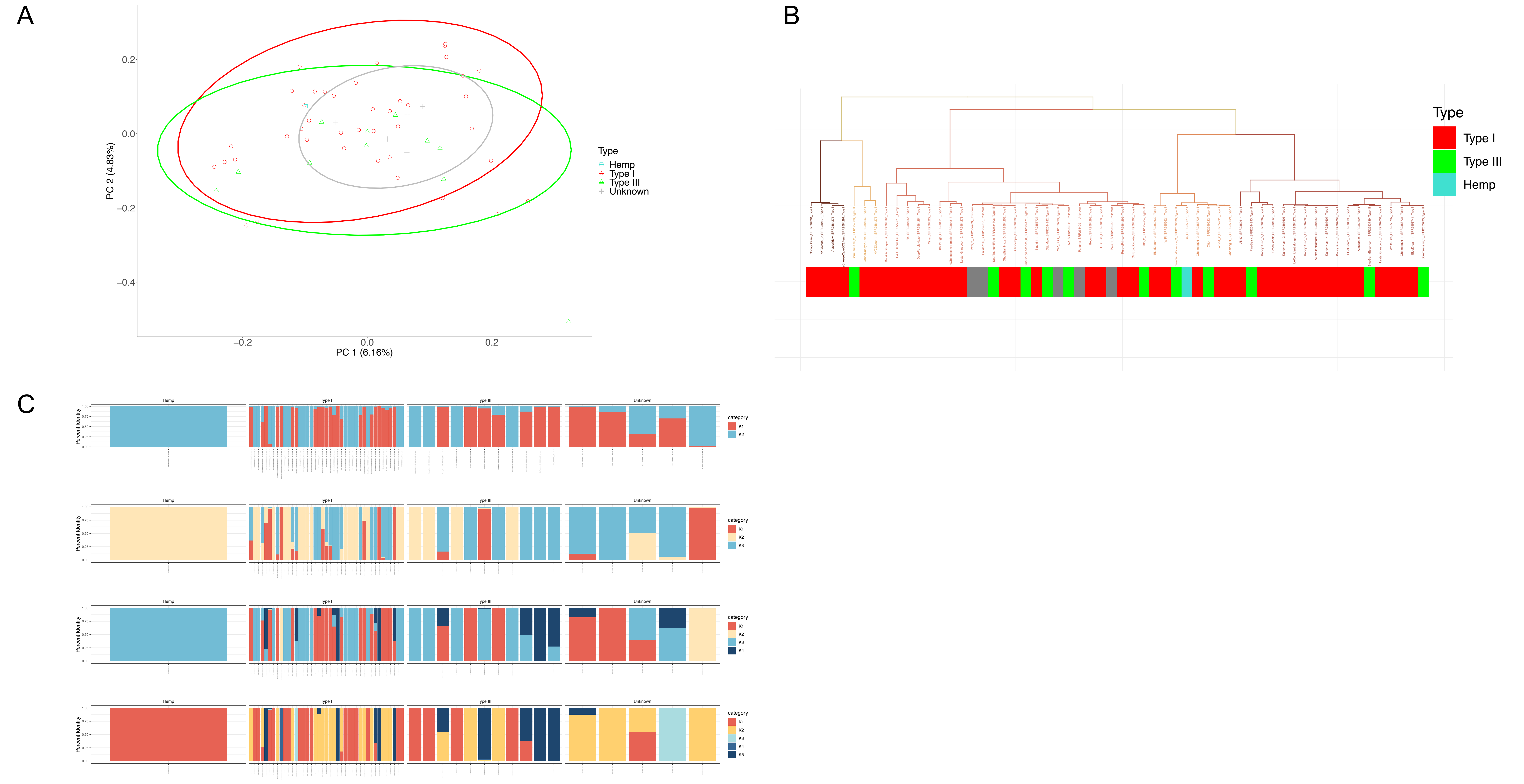


**Supplemental Figure 5** Nuclear SNP analysis for the Courtagen Life Sciences dataset for 58 samples. Clustering was conducted based on nuclear genetic SNPs while reported use-type within the dataset is below in solid bars to facilitate interpretation based upon community standards **(A)** PCA by use-type based on 119 nuclear SNPs. Use-type associations include Hemp (n=1), THC-Dominant (Type I) (n=41), CBD-Dominant (Type III) (n=11) and Unknown (n=5) **(B)** Hierarchical cluster dendrogram with use-type indicated below **(C)** Visualization of population structure and admixture using the fastSTRUCTURE software (k=2-5).

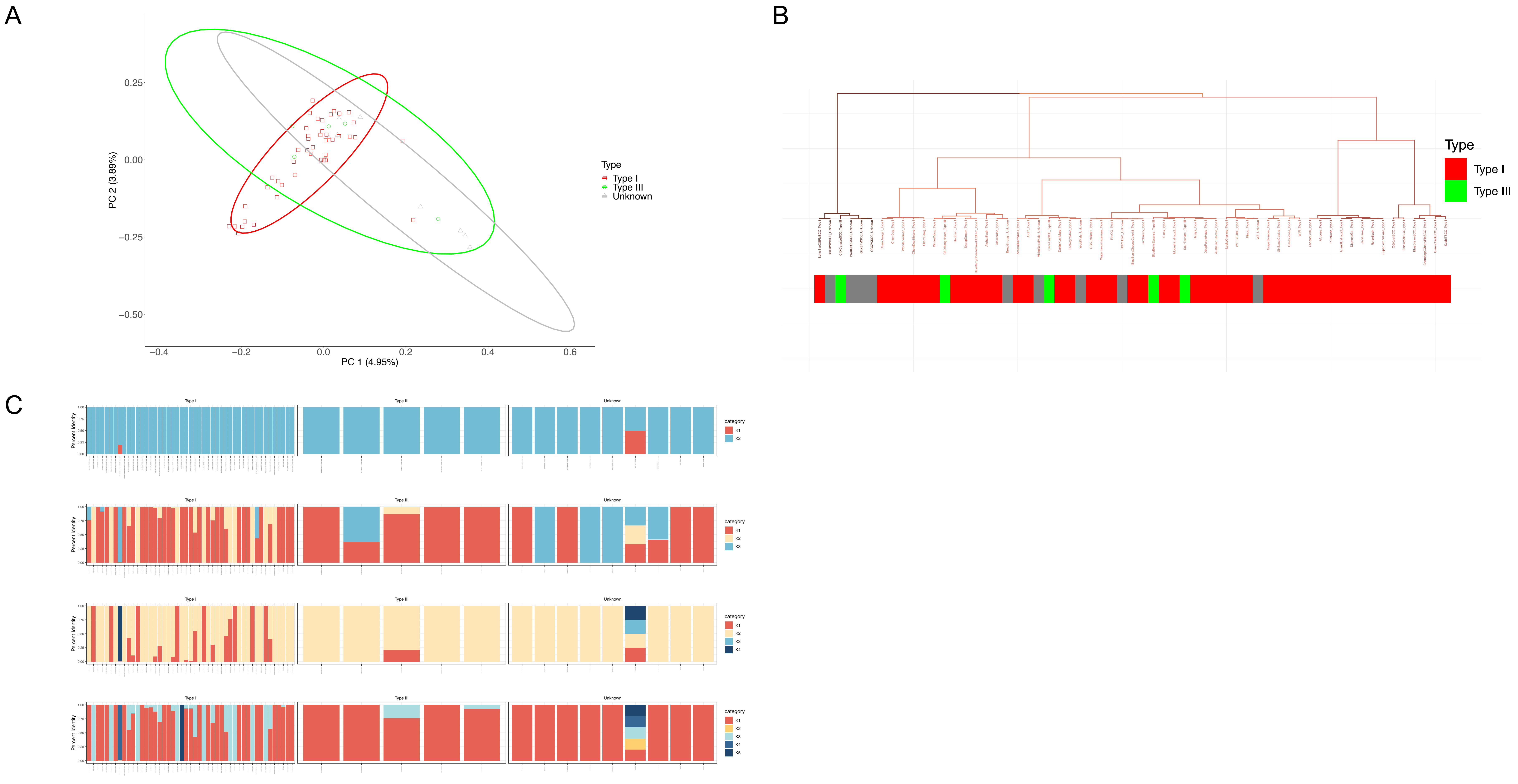


**Supplemental Figure 6** Nuclear SNP analysis for the Medicinal Genomics 61 dataset for 61 samples. Clustering was conducted based on nuclear genetic SNPs while reported use-type within the dataset is below in solid bars to facilitate interpretation based upon community standards **(A)** PCA by use-type based on 2,267 nuclear SNPs. Use-type associations include Hemp (n=1), THC-Dominant (Type I) (n=47), CBD-Dominant (Type III) (n=5) and Unknown (n=9) **(B)** Hierarchical cluster dendrogram with use-type indicated below **(C)** Visualization of population structure and admixture using the fastSTRUCTURE software (k=2-5) with the optimal number of K being 3 using the silhouette method.

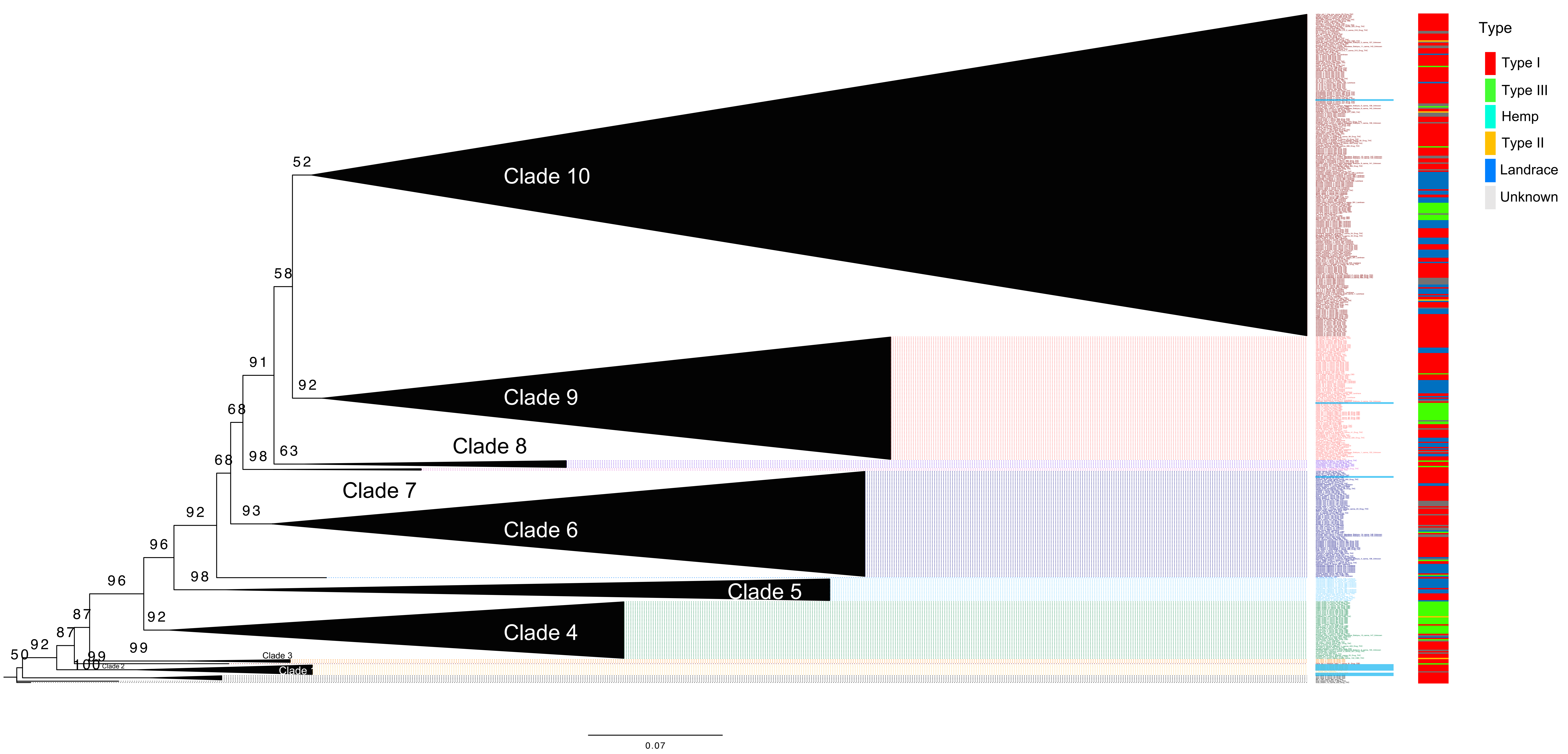


**Supplemental Figure 7** Maximum Likelihood tree for the LeafWorks Inc. dataset constructed from 1,405 nuclear SNPs from 498 samples. Modeltest-ng revealed the TIM2+G4 as the best fit substitution model and IQ-Tree software was used for phylogenetic inference. Blue Dream samples (n=12) are highlighted in blue at the branch tips. Use-type for individual samples is additionally indicated.

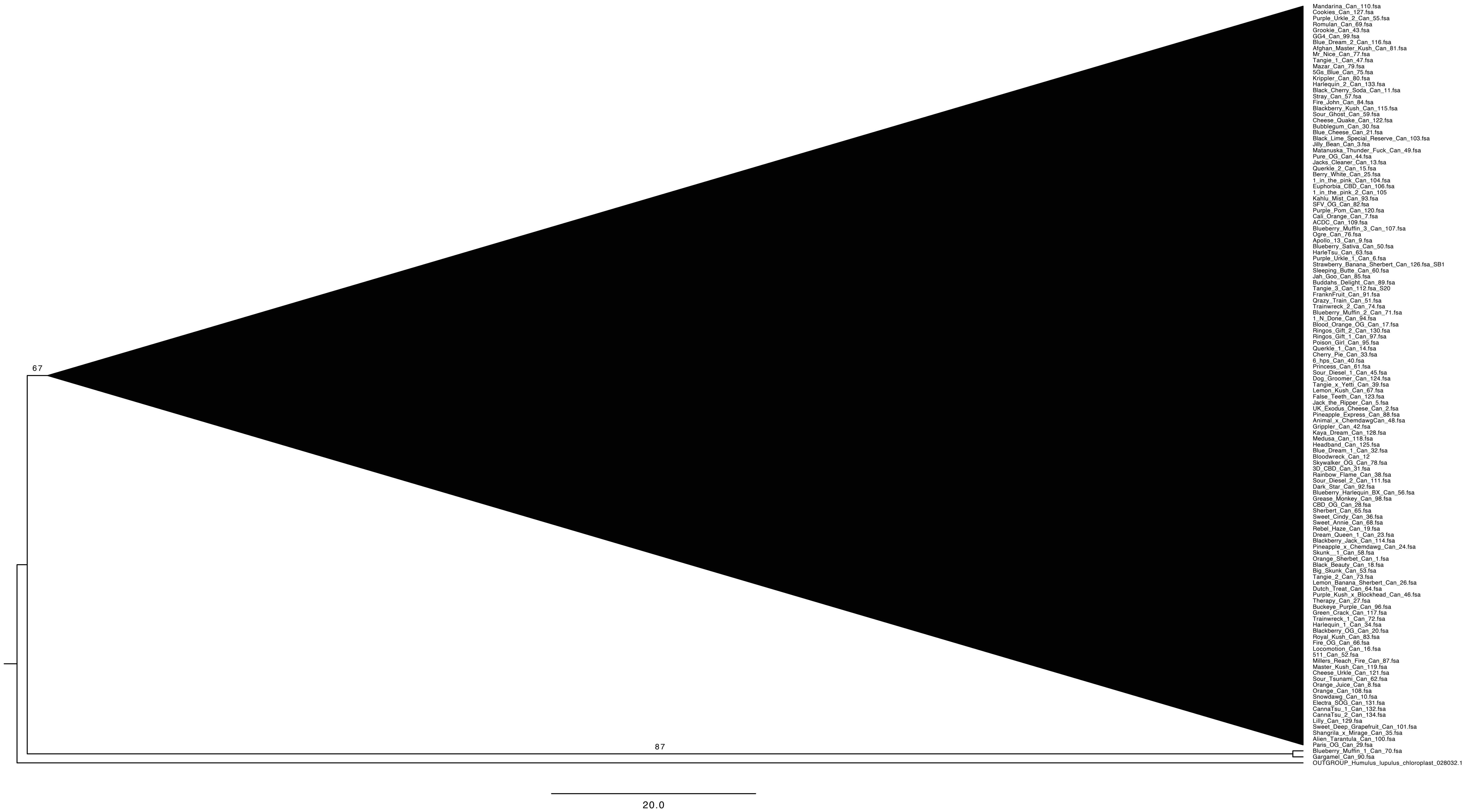


**Supplemental Figure 8** Maximum Likelihood phylogenetic tree for 126 whole chloroplast assemblies. Individuals were aligned using MAFFT. Modeltest-NG revealed the GTR+G4 as the best fit substitution model and IQ-Tree software was used for phylogenetic inference. The resultant tree was visualized using FigTree (Version 1.4.4).

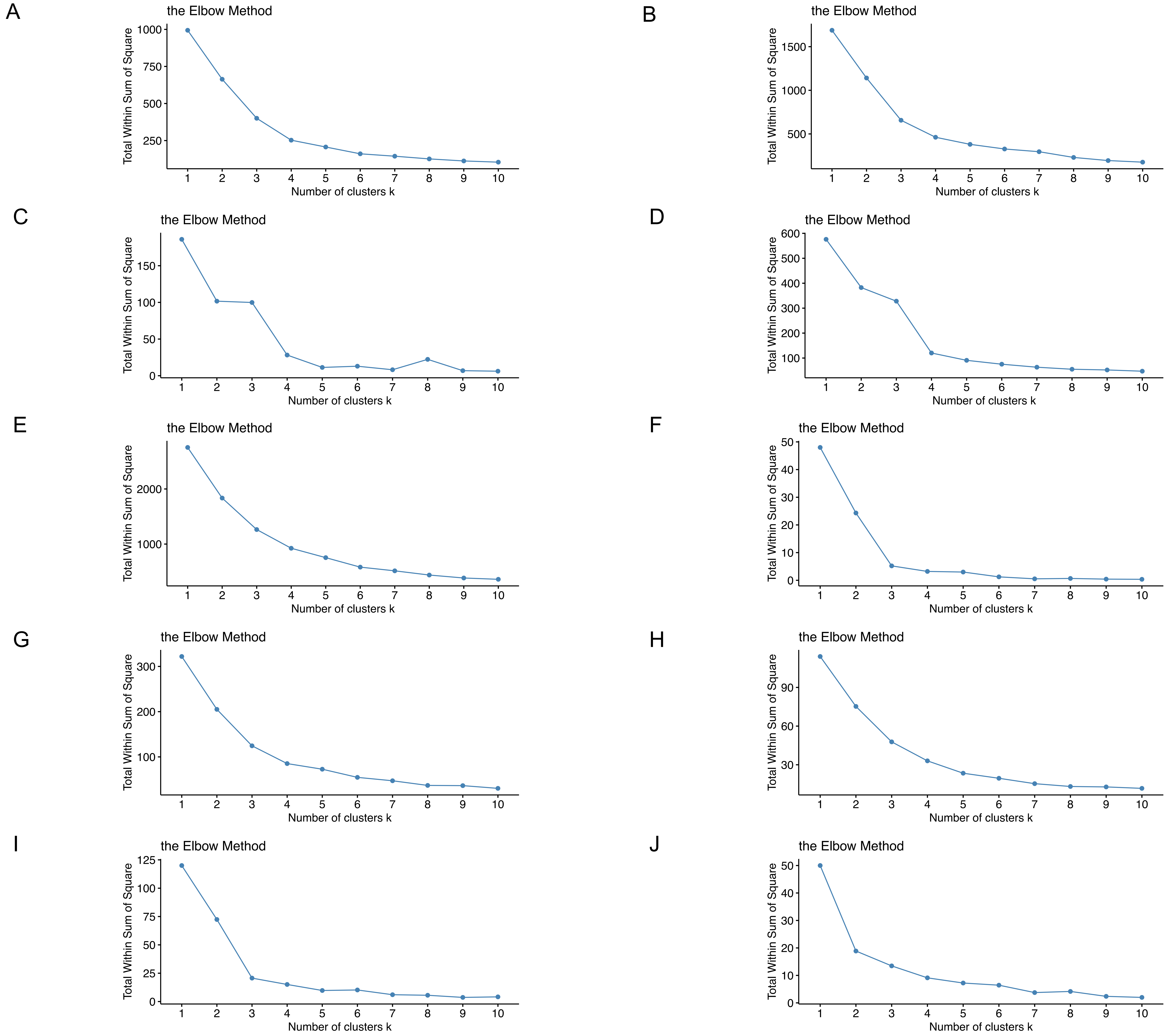


**Supplemental Figure 9** Examining optimal K number across the datasets using the Elbow Method **(A)** LeafWorks Inc. dataset **(B)** Phylos Biosciences dataset (n=845) **(C)** Soorni dataset (n=94) **(D)** Medicinal Genomics StrainSEEK V1 (n=289) **(E)** Phylos Biosciences dataset (n=1378) **(F)** Sunrise Genetics (n=25) **(G)** Colorado dataset (n=162) **(H)** Courtagen dataset (n=58) **(I)** Kannapedia 61 dataset (n=61) **(J)** LeafWorks Inc. landrace samples (n=14).

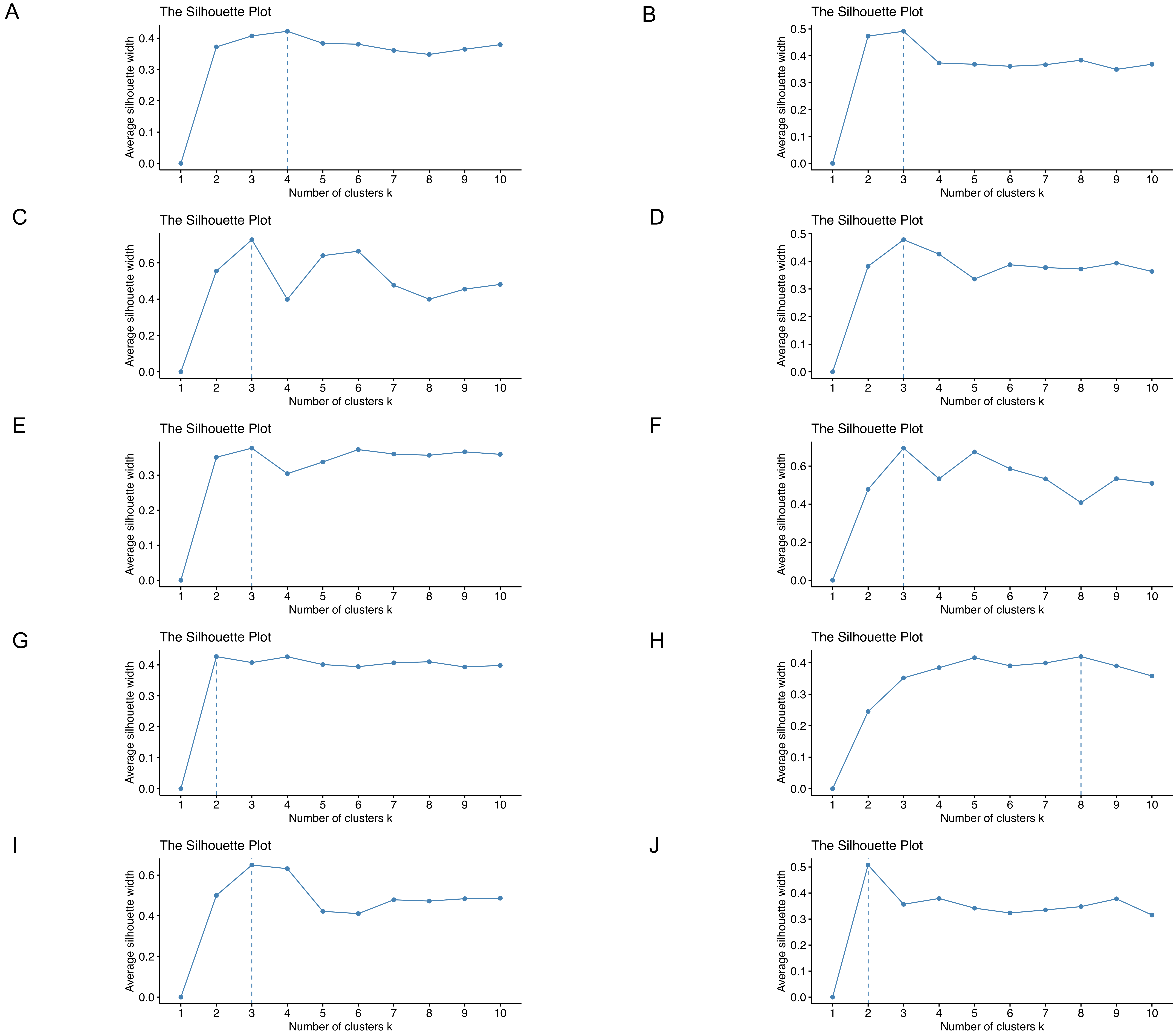


**Supplemental Figure 10** Examining optimal K number across the datasets using the Silhouette Method **(A)** LeafWorks Inc. dataset **(B)** Phylos Biosciences dataset (n=845) **(C)** Soorni dataset (n=94) **(D)** Medicinal Genomics StrainSEEK V1 (n=289) **(E)** Phylos Biosciences dataset (n=1378) **(F)** Sunrise Genetics (n=25) **(G)** Colorado dataset (n=162) **(H)** Courtagen dataset (n=58) **(I)** Kannapedia 61 dataset (n=61) **(J)** LeafWorks Inc. landrace samples (n=14).
